# The molecular basis of coupling between poly(A)-tail length and translational efficiency

**DOI:** 10.1101/2021.01.18.427055

**Authors:** Kehui Xiang, David P. Bartel

## Abstract

In animal oocytes and early embryos, mRNA poly(A)-tail length strongly influences translational efficiency (TE), but later in development this coupling between tail length and TE disappears. Here, we elucidate how this coupling is first established and why it disappears. Overexpressing cytoplasmic poly(A)-binding protein (PABPC) in frog oocytes specifically improved translation of short-tailed mRNAs, thereby diminishing coupling between tail length and TE. Thus, coupling requires limiting PABPC, implying that in coupled systems longer-tail mRNAs better compete for limiting PABPC. In addition to expressing excess PABPC, post-embryonic cells had two other properties that prevented strong coupling: terminal-uridylation-dependent destabilization of mRNAs lacking bound PABPC, and a regulatory regime wherein PABPC contributes minimally to TE. Thus, these results revealed three fundamental mechanistic requirements for coupling and defined the context-dependent functions for PABPC, in which this protein promotes TE but not mRNA stability in coupled systems and mRNA stability but not TE in uncoupled systems.

## Introduction

Most eukaryotic mRNAs are polyadenylated at their 3′ ends in a process associated with transcriptional termination. In the nucleus, these poly(A) tails can facilitate mRNA nucleocytoplasmic export (Kühn and Wahle, 2004), whereas in the cytoplasm, they serve as molecular timers for mRNA decay, with their lengths becoming progressively shorter by deadenylation, which eventually leads to mRNA de-capping and turnover (Goldstrohm and Wickens, 2008; Chen and Shyu, 2011; Eisen et al., 2020). The length of a poly(A) tail can also influence mRNA translational efficiency (TE). Pioneering studies in maturing oocytes and early embryos show that lengthening of poly(A) tails through cytoplasmic polyadenylation is critical for regulating gene expression during these early stages of animal development (Sallés et al., 1994; Sheets et al., 1995; Richter, 1999). Results from these and other single-gene studies in oocytes and early embryos had led to the notion that the length of a poly(A) tail generally correlates with TE (Eckmann et al., 2011; Weill et al., 2012). More recent transcriptome-wide studies confirm a strong global relationship between tail length and TE in oocytes and early embryos (Subtelny et al., 2014; Eichhorn et al., 2016; Lim et al., 2016). However, in fish, frogs, and flies, this correlation diminishes near the time of gastrulation, and coupling between poly(A)-tail length and TE is essentially nonexistent in post-embryonic systems (Subtelny et al., 2014; Eichhorn et al., 2016; Park et al., 2016). Thus, these global analyses reveal a developmental transition in how translation is regulated (Subtelny et al., 2014), which closely follows the long-known maternal-to-zygotic transition in transcriptional control. The existence of this transition in translational control brings to the fore mechanistic questions as to how coupling between poly(A)-tail length and TE is established in oocytes and early embryos and why this coupling disappears later in development.

Cytoplasmic poly(A)-binding proteins (PABPCs) are highly conserved RNA-binding proteins in eukaryotes (Mangus et al., 2003). Although *Saccharomyces cerevisiae* has only one PABPC (Pab1p), most animals contain multiple paralogs that have spatially and temporally varied expression patterns (Smith et al., 2014; Wigington et al., 2014). PABPCs have high affinity to poly(A) sequences in vitro (*K*_d_ ∼5 nM for A_25_) and require at least 12 As for efficient binding (Kühn and Wahle, 2004). Binding of PABPCs to mRNA poly(A) tails can enhance translation, but the mechanism of this enhancement is unclear. One model posits that the mRNA forms a closed-loop structure mediated by the association of the eukaryotic translation initiation factor eIF4G (a scaffolding protein) with both PABPC and the cap-binding protein eIF4E (Wells et al., 1998; Hinnebusch, 2014; Thompson and Gilbert, 2016). This association is proposed to stabilize the interaction between eIF4E and the mRNA 5′ cap and facilitate recruitment and/or recycling of ribosomes to increase translation initiation (Kahvejian et al., 2001). However, despite direct visualization of loop-like assemblies both within some cells and in an in vitro reconstituted system (Christensen et al., 1987; Wells et al., 1998), results of several studies have questioned the universality of this model among different mRNAs and biological systems (Amrani et al., 2008; Costello et al., 2015; Thompson and Gilbert, 2016; Rissland et al., 2017; Adivarahan et al., 2018).

PABPCs can also influence mRNA stability, as shown in yeast. Genetic ablation of yeast Pab1p is lethal and causes lengthening of steady-state poly(A)-tail lengths (Sachs and Davis, 1989), which is attributed to pre-mature mRNA decapping and compromised deadenylation (Caponigro and Parker, 1995). Both yeast and mammalian PABPCs can interact with two mRNA deadenylation complexes PAN2-PAN3 and CCR4-NOT, and either promote or inhibit their activities in vitro (Uchida et al., 2004; Webster et al., 2018; Yi et al., 2018; Schäfer et al., 2019). Because mRNA decay is coupled to deadenylation (Decker and Parker, 1993; Eisen et al., 2020), the deadenylation-stimulatory effects of PABPC would accelerate the demise of bound mRNAs, which contrasts to other studies suggesting PABPC protects mRNAs from degradation in cell extracts (Bernstein et al., 1989; Wang et al., 1999). The dichotomous and potentially conflicting functions of metazoan PABPC examined in vitro raise the question of the extents to which PABPC might influence mRNA poly(A)-tail length and stability in metazoan cells.

PABPCs are generally thought to coat mRNA poly(A) tails in the cytoplasm (Kühn and Wahle, 2004; Lima et al., 2017). However, the stoichiometry between PABPC and poly(A) sites might vary in different biological contexts (Voeltz et al., 2001; Cosson et al., 2002), and it is unclear whether this potentially variable stoichiometry might impact gene regulation in cells. Moreover, the possibility that PABPC might influence protein synthesis by affecting either mRNA stability or TE can complicate analysis of its molecular functions in different biological systems, leaving its mechanistic roles poorly understood.

Here, we uncover mechanistic requirements for coupling between poly(A)-tail length and TE observed in oocytes and early embryos, showing that this coupling and the subsequent uncoupling observed later in development rely on a context-dependent switch in the function of PABPCs.

## Results

### Limiting PABPC is required for tail length to strongly influence TE of reporter mRNAs

To assay the influence of poly(A)-tail length on TE, we used an in vitro translation extract made from stage VI *Xenopus laevis* oocytes, where cytoplasmic polyadenylation leads to translational activation of the *c-mos*, *cdk2* and some cyclin mRNAs (Stebbins-Boaz and Richter, 1994; Sheets et al., 1995). Into this extract we added *Renilla* luciferase (*Rluc*) reporter mRNAs with either a short (30 nt) or a long (120 nt) poly(A) tail (Figure 1A). These mRNAs were made by in vitro transcription from DNA templates that encoded the mRNA body followed by the poly(A) tail as well as the hepatitis delta virus (HDV) self-cleaving ribozyme, which cleaved during in vitro transcription to generate not only a defined 3′ end at the desired poly(A)-tail length but also a 2′-3′-cyclic phosphate designed to inhibit undesired lengthening or shortening of the tail (Avis et al., 2012).

**Figure 1.**
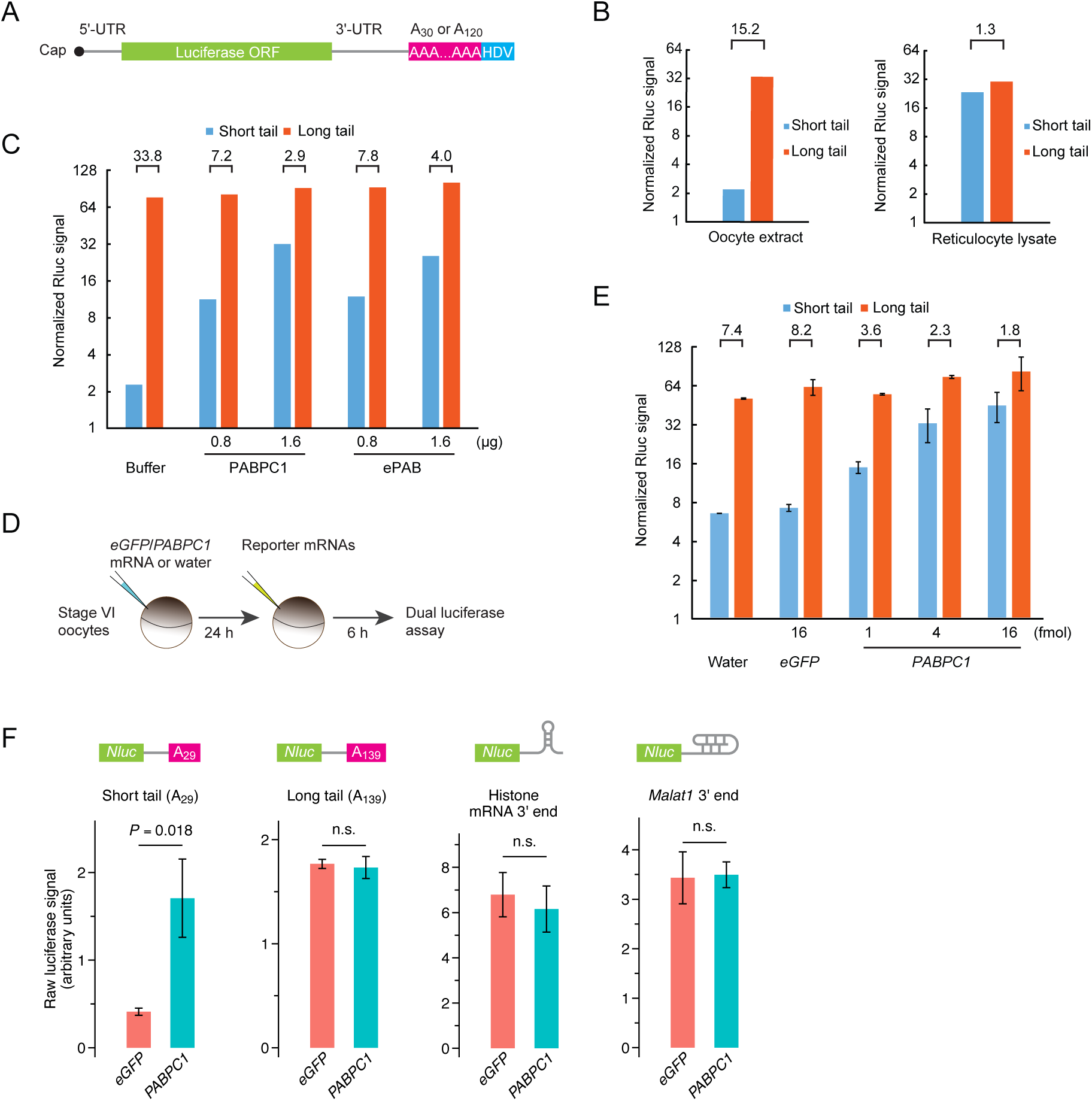
PABPC overexpression uncouples p(A)-tail length and TE in frog oocytes. (**A**) Schematic of capped T7 transcripts with two different tail lengths, which were used as reporter mRNAs. Additional sequences beyond the HDV sequence are not shown. (**B**) The effect of tail length on relative yields of in vitro translation of short- and long-tailed reporter mRNAs, in either frog oocyte extract (left) or rabbit reticulocyte lysate (right). The number above each bracket indicates the fold difference. (**C**) The effect of purified PABPC on relative yields of in vitro translation of short- and long-tailed reporter mRNAs in frog oocyte extract. Purified PABPC1 and ePAB were each added as indicated. Otherwise, this panel is as in (**B**). (**D**) Experimental scheme for serial-injection of mRNAs into frog oocytes. (**E**) The effect of PAPBC1 overexpression on relative translation of reporter mRNAs into frog oocytes. Differential PABPC1 expression was achieved by injecting indicated amount of mRNA in the first injection (error bars, standard deviations from two biological replicates). Otherwise, this panel is as in (**B**). (**F**) The effect of PAPBC1 overexpression on translation of reporter mRNAs with different 3′-end structures in frog oocytes. Shown are raw luciferase yields from *NanoLuc* reporters (*Nluc*) that have either a short poly(A) tail, a long poly(A) tail, a histone mRNA 3′-end stem-loop, or a *Malat1* triple-helix 3′-end in oocytes overexpressing either eGFP or PABPC1 (error bars, standard deviations from three biological replicates). *P* values are from one-sided *t*-tests (n.s., not significant). The following figure supplement is available for figure 1: **Figure supplement 1**. Supporting data for reporter experiments examining the effect of PAPBC levels on coupling between tail length and translation.

When added to the frog oocyte extracts together with a firefly luciferase (*Fluc*) mRNA, used to normalize for overall translation activity, the long-tailed reporter was translated substantially better than was the short-tailed reporter (Figure 1B). In contrast, the same reporter mRNAs were translated nearly equally well in rabbit reticulocyte lysate, a post-embryonic differentiated system for which no coupling between tail length and TE was expected (Subtelny et al., 2014) (Figure 1B). In both systems, the reporter mRNAs were stable with no detectable changes to their tail lengths (Figure 1—figure supplement 1A–C). Thus, the large difference in luciferase signal observed between the short- and long-tailed reporters in the oocyte extract was attributable to a difference in TE. These results showed that the causal relationship between longer poly(A)-tail length and greater TE observed for some maturation-specific mRNAs in frog oocytes (Stebbins-Boaz and Richter, 1994; Sheets et al., 1995) is not unique to those mRNAs, and indicated that frog oocyte extracts provide a system for probing the mechanism that couples tail length to TE.

When considering the potential mechanisms for reading out tail length and promoting translation, a role for PABPC seemed plausible. For instance, translation might be sensitive to the number of PABPC molecules associated with an mRNA. In one mechanistic possibility, PABPC might be in excess over its binding sites within tails, such that tails are coated with the protein, as is generally thought to occur (Mangus et al., 2003; Lima et al., 2017), in which case, mRNAs with longer tails might be detected as those able to bind more PABPC molecules. At another mechanistic extreme, PABPC might be limiting, such that mRNAs compete with each other for PABPC binding, in which case, those with long poly(A)-tail lengths would compete more effectively and thereby preferentially benefit from any enhancement in TE that PABPC binding confers. To distinguish between these possibilities, we increased available PABPC in our oocyte extracts, reasoning that if PABPC were already coating the tails, adding more would have little effect, whereas if PABPC were limiting, adding more would diminish the competition for PABPC binding and thereby reduce the difference in TE observed between short-and long-tailed mRNAs. Accordingly, we purified recombinant *Xenopus* PABPC1 and embryonic PABPC (ePAB), to near homogeneity (Figure 1—figure supplement 1D) and examined their influence when added to the in vitro translation extract derived from stage VI oocytes. As more of either of these proteins was added, the normalized translation signal of the short-tailed reporter approached that of the long-tailed reporter (Figure 1C). This concentration-dependent diminution of coupling between tail-length and TE strongly supported the hypothesis that limiting PAPBC was required for this coupling.

To investigate whether this requirement of limiting PABPC was restricted to our in vitro extracts or whether it also applied to living oocytes, we performed serial-injection experiments in oocytes. Stage VI frog oocytes were first injected with either *PABPC1* mRNA or a control, and after waiting 24 h to allow PABPC1 protein to accumulate, oocytes were injected with the reporter mRNAs and assayed for luciferase activity (Figure 1D, Figure 1—figure supplement 1E). Whereas, injecting the control mRNA, *eGFP*, had no more influence than injecting water, injecting *PABPC1* mRNA significantly reduced the extent to which poly(A)-tail length and TE were coupled, and it did so in a concentration-dependent manner (Figure 1E). Reporter poly(A)-tail lengths did not change over the course of the experiment (Figure 1–figure supplement 1F), which indicated that the increased relative translation of the short-tailed reporter mRNA was not due to elongated poly(A) tails. Similar results were observed when either injecting *ePAB* mRNA rather than *PABPC1* mRNA (Figure 1— figure supplement 1G), replacing *Rluc* mRNA reporters with analogous *NanoLuc* (*Nluc*) reporters (Figure S1H), or injecting purified PABPC1 protein rather than *PABPC1* mRNA (Figure 1—figure supplement 1I).

Introducing additional PABPC into frog oocytes specifically improved translation of the short-tailed reporter while having little effect on translation of either the long-tailed reporter or reporters for which tails were replaced with either a stem-loop from the 3′ end of a histone mRNA (Ling et al., 2002) or a triple-helix from the 3′ end of the *Malat1* non-coding RNA (Wilusz et al., 2012) (Figure 1F). The observation that mRNAs required a tail to benefit from added PABPC indicated that the effects of adding PABPC were direct and not some secondary consequence of altering translation. Moreover, the observation that PABPC had little effect on translation of long-tailed mRNAs suggested that these mRNAs more effectively competed for the limiting PABPC and thus were efficiently translated even in the absence of added PABPC. This negligible effect on translation of long-tailed mRNAs also implied that just a few bound PABPC molecules (perhaps only one) might suffice to boost translation, with little added benefit of additional PABPC binding, which disfavored a coupling mechanism that simply counts the number of bound PABPC molecules.

In summary, our results with reporters in oocytes and oocyte extracts confirmed both the positive effect of PABPC on translation and the causal relationship between poly(A)-tail length and TE in these systems. Moreover, these results implied that TE is not simply a function of the number of PABPC molecules coating the tail and revealed that strong coupling between poly(A)-tail length and TE requires limiting PABPC.

### Limiting PABPC is required for tail length to strongly influence TE of endogenous mRNAs

To examine the global effect of increasing PABPC on the translational regulatory regime acting in the oocyte, we monitored the relationship between tail length and TE for endogenous mRNAs of the oocytes. As expected from results of single-gene experiments in frog oocytes (Figure 1E) (Stebbins-Boaz and Richter, 1994; Sheets et al., 1995) and the strong coupling between poly(A)-tail length and TE observed in both frog embryos and fly oocytes (Subtelny et al., 2014; Eichhorn et al., 2016; Lim et al., 2016), we found that poly(A)-tail length correlated strongly with TE in stage VI frog oocytes (Figure 2A). Overexpressing either PABPC1 or ePAB in these oocytes significantly diminished the coupling, with the Spearman correlation (*R*_s_) for the relationship between tail length and TE dropping from 0.62 to 0.36 and 0.38, respectively (Figure 2A, both *P* = 0, modified Dunn and Clark’s *z*-test (Diedenhofen and Musch, 2015)). In contrast, overexpressing eGFP had no significant impact on the coupling (*P* = 0.11), which showed that this transcriptome-wide effect was a result of additional PABPC protein rather than a non-specific effect of adding more mRNA.

**Figure 2.**
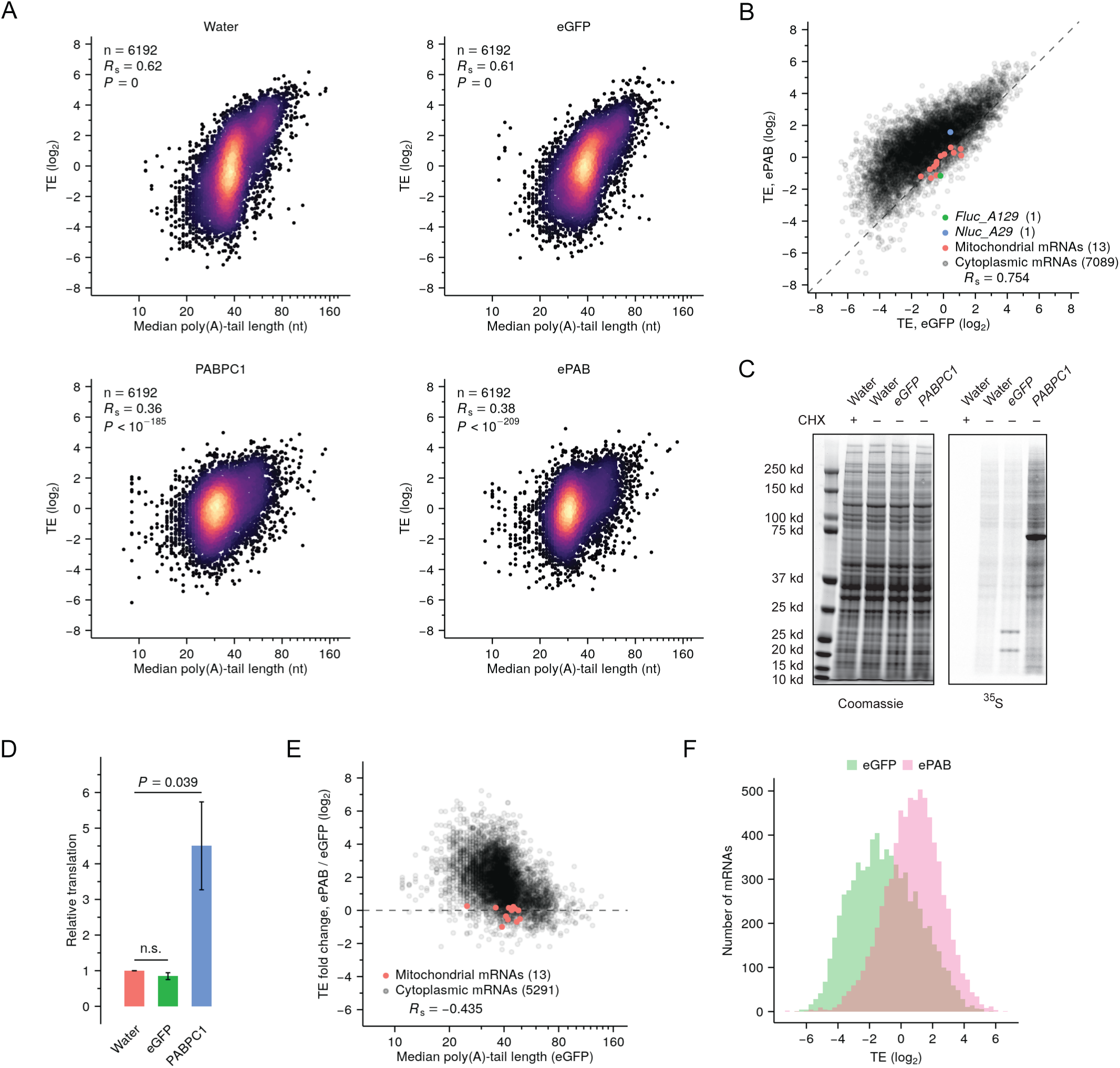
Increased PABPC promotes translation of endogenous short-tailed mRNAs, thereby diminishing coupling between tail length and TE. (**A**) The effect of PABPC on coupling between tail length and TE in frog oocytes. Shown is the relationship between TE and median poly(A)-tail length in oocytes injected with either water or mRNAs encoding either eGFP, PABPC1, or ePAB. Results are shown for mRNAs from genes with ≥ 100 poly(A) tags. *R*_s_ is the Spearman’s correlation coefficient. (**B**) A global increase in TE observed upon overexpressing ePAB. TE in ePAB-overexpressing oocytes is compared with TE in eGFP-expressing oocytes. Experiment as in A, except rRNAs were not depleted. Also shown are results from co-injected a short-tailed *Nluc* reporter and a long-tailed *Fluc* reporter. (**C**) Effect of expressing PABPC1 on protein synthesis in oocytes. At the left is a gel showing total protein after injection of the indicated mRNAs (or water) or treatment with cycloheximide (CHX), as visualized by Coomassie staining. At the right is the same gel showing protein synthesis, as visualized by incorporation of ^35^S-methionine and ^35^S-cysteine. (**D**) Quantification of the effect of expressing PABPC1 on protein synthesis in oocytes, as measured in (**C**) and two additional biological replicates. Only regions above the PABPC1 band, and between the PABPC1 band and the top eGFP band were used for quantification. Values were normalized to that of the mean value from water-injected oocytes (error bars, standard deviation; *P* values, one-sided *t*-tests; n.s., not significant). (**E**) The preferential effect of overexpressing ePAB on the TE of short-tailed mRNAs. TE fold changes observed between ePAB-overexpressing and eGFP-expressing oocytes are plotted as a function of median tail length in eGFP-expressing oocytes. TE and tail-length values were obtained from different batches of oocytes; results are shown for mRNAs from genes with ≥ 100 poly(A) tags. (**F**) Effect of overexpressing ePAB on the distribution of TE values observed in frog oocytes. Shown is the TE distribution observed in ePAB-overexpressing oocytes and that observed in eGFP-expressing oocytes. The following figure supplement is available for figure 2: **Figure supplement 2**. Increased PABPC increases translation of endogenous short-tailed mRNAs in frog oocytes.

Accompanying the reduced coupling observed upon PABPC overexpression was a significant relative increase of TE for short-tailed mRNAs, an effect not observed in eGFP-expressing oocytes (Figure 2—figure supplement 2A). This TE increase was not accompanied by corresponding lengthening of poly(A) tails (Figure 2—figure supplement 2B), implying that tail-length changes did not cause relative TE changes. To make comparisons of absolute TE changes, we repeated the ePAB-overexpression experiment but omitted rRNA depletion during sequencing library construction, thereby allowing us to normalize TE using mitochondrial mRNAs (Iwasaki et al., 2016), which were otherwise depleted by Illumina Ribo-Zero kits (Figure 2—figure supplement 2C). In this experiment, we also injected oocytes with a short-tailed *Nluc* mRNA reporter and a long-tailed *Fluc* mRNA reporter and monitored their absolute TE changes together with those of endogenous mRNAs. Most endogenous mRNAs had greater absolute TE in ePAB-overexpressing oocytes compared to eGFP-expressing control oocytes (Figure 2B). This result was consistent with ^35^S metabolic-labeling experiments showing that overexpression of PABPC1 but not eGFP significantly increased global protein synthesis in oocytes (Figure 2C–D). Moreover, the magnitude of the TE increase conferred by ePAB-overexpression negatively correlated with tail length, which showed that translation of short-tailed mRNAs improved substantially more than that of long-tailed mRNAs (Figure 2E). Indeed, adding ePAB had essentially no overall effect on TE of endogenous mRNAs with the longest tails (median TE fold change = 1.06 for the 54 mRNAs with median tail lengths > 80 nt), which further supported the notion that initial PABPC binding is more consequential than subsequent binding. The preferential improvement of TEs for short-tailed mRNAs led to not only an overall shift in TE but also narrowing of the TE distribution (Figure 2F) to more closely resemble the distributions observed in cells in which poly(A)-tail length and TE are not coupled (Subtelny et al., 2014). These results supported the hypothesis that increasing PABPC in oocytes increases the opportunity for short-tailed mRNAs to bind a PABPC molecule, thereby promoting translation.

Overall, the results of our global analyses of mRNAs in frog oocytes agreed with those of reporter assays, thereby extending to endogenous mRNAs support for the conclusion that TE is not simply a function of the number of PABPC molecules coating each poly(A) tail. Instead, limiting PABPC plays a critical role in conferring strong coupling between poly(A)-tail length and TE.

### Intragenic analyses further demonstrate the importance of limiting PABPC for establishing coupling between tail length and TE

Our global analysis examining the relationship between poly(A)-tail length and TE of endogenous mRNAs in oocytes differed from our reporter assays in that the comparison was made between mRNAs of different genes, which can be confounded by features other than tail length that vary between these mRNAs. To overcome this issue, we developed a high-throughput method for comparing effects on different tail-length isoforms from each gene. This approach for intragenic analyses, called PAL-TRAP (<u>P</u>oly(<u>A</u>) tail-<u>L</u>ength profiling following <u>T</u>ranslating <u>R</u>ibosome <u>A</u>ffinity <u>P</u>urification), resembled other TRAP approaches in that ribosomes were sparsely tagged such that their immunoprecipitation (IP) preferentially isolated mRNA isoforms associated with more ribosomes, which were inferred to be more highly translated (Heiman et al., 2008; Chen and Dickman, 2017). In a system in which poly(A)-tail length and TE were coupled, longer-tail mRNAs were expected to be associated with more ribosomes and therefore enriched in the eluate (Figure 3A), whereas in an uncoupled system, longer-tail mRNAs were not expected to be enriched in the eluate.

**Figure 3.**
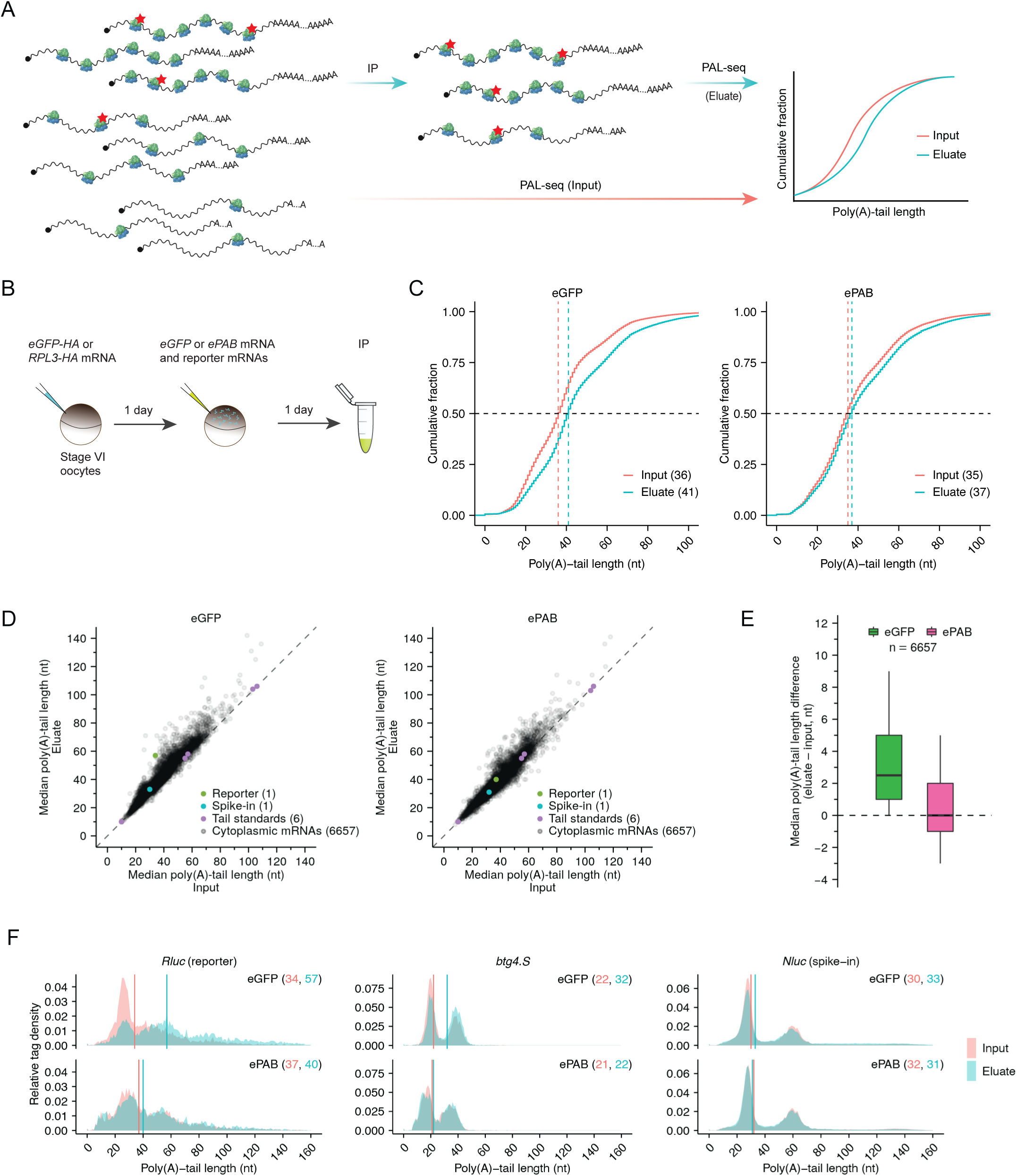
Limiting PABPC is required for intragenic coupling between poly(A)-tail length and TE. (**A**) The PAL-TRAP method for measuring intragenic effects of tail length on TE. Ribosomes are sparsely tagged so that highly translated mRNAs are more likely to contain tagged ribosomes and thus be enriched in the immunoprecipitation (IP) eluate. Tail lengths of both input and eluate mRNAs are measured and compared for mRNAs of each gene. Enrichment of long-tailed isoforms in the eluate indicates that poly(A)-tail length and TE are coupled, whereas no enrichment indicates otherwise. (**B**) The experimental scheme of PAL-TRAP in frog oocytes. See figure supplement 3B–C for results from pulldowns using the eGFP-HA control. (**C**) Effect of overexpressing ePAB on coupling between tail length and TE, as detected after pooling PAL-TRAP results for mRNAs from different genes. Plotted are cumulative distributions of poly(A)-tail lengths in the PAL-TRAP input and eluate obtained after expressing either eGFP (left) or ePAB (right) in oocytes. Median values are indicated (dashed lines) and listed in parentheses. (**D**) The effect of overexpressing ePAB on intragenic coupling between tail length and TE. Plotted for each mRNA isoform is the median poly(A)-tail length of mRNAs in the PAL-TRAP eluate compared to that in the input. Shown are results for mRNAs from oocytes either expressing eGFP or overexpressing ePAB (left and right, respectively). Each point represents an mRNA isoform with a unique 3′ end represented by ≥100 poly(A) tags in both input and eluate. Also indicated are results for 1) an *Rluc* reporter mRNA possessing a variable-length tail (reporter), which was co-injected with mRNAs expressing either eGFP or ePAB, 2) an mRNA with a variable-length tail, which was spiked into the lysate immediately before IP (spike-in), and 3) synthetic RNAs with defined tail lengths added to samples prior to library preparation (tail standards). Points for eight standards with longer tails fell outside the plot areas, as did a point representing the mRNA 3′-end from one gene (*uqcrb.S*) in the ePAB sample. (**E**) Summary of differences in median tail lengths observed between the eluate and the input of mRNA isoforms shown in (**D**). Box and whiskers indicate the 10^th^, 25^th^, 50^th^, 75^th^ and 90^th^ percentiles. (**F**) Effect of over-expressing ePAB on intragenic tail-length distributions in frog oocytes. Shown are tail-length distributions of the reporter *Rluc* (left), an endogenous oocyte mRNA *btg4.S* (middle), and the spike-in mRNA *Nluc* (right) in eGFP-expressing (top) or ePAB-overexpressing (bottom) oocytes. Median tail-length values are indicated (vertical lines) and listed in parentheses. The following figure supplement is available for figure 3: **Figure supplement 3**. Supporting data for PAL-TRAP analyses in frog oocytes.

To implement PAL-TRAP, we first injected stage VI frog oocytes with an mRNA encoding C-terminal HA-tagged RPL3 (Chen and Dickman, 2017) and allowed RPL3-HA protein expression and incorporation into ribosomes (Figure 3B). As a control, HA-tagged eGFP was expressed in separate oocytes. To examine the requirement of limiting PABPC for coupling between poly(A)-tail length and TE, we then injected oocytes with mRNAs that expressed either eGFP or ePAB. Confirming that RPL3-HA was incorporated into functional ribosomes, HA-IP from RPL3-HA-expressing oocytes enriched for proteins from both ribosomal subunits, cytoplasmic mRNAs, and ePAB, whereas HA-IP from eGFP-HA-expressing oocytes did not (Figure 3—figure supplement 3A–C). In control oocytes expressing eGFP, longer-tail mRNAs were enriched in the eluate compared to the input (median tail lengths 41 and 36 nt, respectively), which reflect the coupling between tail length and TE (Figure 3C). Overexpression of ePAB reduced this enrichment for longer-tail mRNAs in the eluate compared to the input (median tail lengths 37 and 35 nt, respectively), as expected if limiting PABPC was required for this coupling (Figure 3C). We also analyzed the flowthrough fractions from these pulldown experiments. For both eGFP- and ePAB-expressing oocytes, the cumulative distribution of poly(A)-tail lengths from the flowthrough was nearly identical to that from the input (Figure 3— figure supplement 3D), as expected when considering that only a small fraction of input was depleted by HA-beads. These analyses indicated that our methods were able to capture small differences in tail-length distributions.

When analyzing, for each gene, mRNAs with different tail-length isoforms, mRNAs from most genes (84.4%) had longer median poly(A)-tail lengths in the eluate than in the input (Figure 3D–E), implying that for mRNAs from most genes, long-tailed isoforms were translated more efficiently than short-tailed ones. Although the median of the median tail-length differences was moderate (+2.5 nt), 28.3% mRNAs had a ≥ 5 nt longer median tail length in the eluate (Figure 3E). In contrast, when input and flowthrough were compared, the median tail-length differences centered on 0 nt, and mRNAs from only 5.4% of genes had a ≥ 5 nt longer median tail length in one of the samples, as expected when considering that only a small fraction of input was depleted by HA-beads (Figure 3—figure supplement 3E–F). Overexpressing ePAB reduced the number of genes with long-tailed isoforms enriched in the eluate and shifted the distribution of median tail-length differences closer to 0 nt (Figure 3D–E), as expected if coupling between poly(A)-tail length and TE diminished.

In these PAL-TRAP experiments, a mixture of *Rluc* reporter mRNA molecules with different poly(A)-tail lengths was co-injected with mRNAs that expressed either eGFP or ePAB. In eGFP-expressing oocytes, longer-tail *Rluc* isoforms were highly enriched in the eluate compared to the input, whereas in ePAB-overexpressing oocytes, this difference diminished dramatically (Figure 3F). As expected, *Nluc* reporter mRNA, which was added to the lysate as a spike-in during HA pulldown, was not significantly enriched in longer-tail species in the eluate, regardless of the treatment, which indicated that the changes observed for *Rluc* mRNA reflected changes occurring in the oocyte, prior to lysis (Figure 3F). Although endogenous mRNAs from some genes, such as *btg4.S*, also underwent large changes in median tail differences in response to ePAB-overexpression (Figure 3F), changes were typically smaller for endogenous mRNAs (Figure 3E), which was at least partly attributable to the narrower range of initial tail length isoforms for endogenous mRNAs compared to the injected *Rluc* mRNA.

Taken together, the PAL-TRAP results revealed intragenic coupling between poly(A)-tail length and TE for endogenous mRNAs as well as reporters in stage VI frog oocytes. Moreover, these PAL-TRAP analyses showed that as observed both in our reporter assays and in our global intergenic comparisons, substantial coupling between tail length and TE requires limiting PABPC.

### Additional conditions besides limiting PABPC are required for strong coupling

Having established the necessity of limiting PABPC for coupling, we investigated if it could also be sufficient, i.e., whether limiting PABPC could confer coupling between poly(A)-tail length and TE in cells in which these features were normally uncoupled. To this end, we knocked down PABPCs in HeLa cells. Based on available mRNA-seq and mass spectrometry results (Nagaraj et al., 2011), PABPC1 and PABPC4 are the two major PABPC paralogs in HeLa cells, with PABPC1 four times more abundant than PABPC4 and the two together accounting for > 95% of all PABPC in HeLa cells. Consistent with the idea that PABPC is not normally limiting in uncoupled systems, PABPC1 is estimated to be present in 3-fold excess over the poly(A) sites in HeLa cells (Görlach et al., 1994). To reduce PABPC to limiting levels, we used siRNAs to reduce either PABPC1 or PABPC4 alone or both PABPC1 and PABPC4 by > 90% (Figure 4—figure supplement 4A) and examined the relationship between median tail lengths and TE. Knocking down PABPC4 alone had little impact on the coupling (Figure 4—figure supplement 4B), consistent with the inference that PABPC4 constitutes less than 20% of the total PABPC protein. Although correlation between median poly(A)-tail length and TE gradually increased as more PABPC was depleted, it reached an *R*_s_ of only 0.18 (*P* < 10^−19^) in double-knockdown cells (Figure 4—figure supplement 4B), which was much weaker than that observed in frog oocytes (Figure 2A) and frog and fish early embryos (Subtelny et al., 2014). Minor *R*_s_ increases were observed in other mouse and human post-embryonic cell lines in which PABPC was depleted, but strong coupling between poly(A)-tail length and TE was not established, with no *R*_s_ values exceeding 0.3 (data not shown). Thus, other conditions in addition to limiting PABPC must also be met to confer strong coupling between poly(A)-length and TE.

### PABPC depletion causes premature decay of short-tailed mRNAs in mammalian cells

A striking consequence of depleting PABPC in HeLa cells was a sharp increase in median poly(A)-tail lengths, which for HeLa mRNAs increased an average of 17 and 39 nt in PABPC1- and double-knockdown cells, respectively (Figure 4A). Substantial changes were also observed in the distributions of global poly(A)-tail length, which showed that mRNAs with tails ranging from 10–50 nt were > 2-fold depleted in the PABPC1-knockdown cells and that mRNAs with tails ranging from 10–135 nt were 2- to 20-fold depleted in the double-knockdown cells (Figure 4B). Similar results were obtained in NIH3T3 cells (Figure 4—figure supplement 5A–B). In contrast, loss of short-tailed mRNAs was not observed for mitochondria-encoded mRNAs, as expected for these mRNAs that never encounter PABPC (Figure 4A–B, Figure 4—figure supplement 5A–B), and examination of internal standards and replicates confirmed that the loss of short-tailed cytoplasmic mRNAs was not attributable to inaccurate or variable measurements (Figure 4—figure supplement 5C–D). Moreover, knocking down the minor isoform (PABPC4) alone did not have a similar effect (Figure 4—figure supplement 5E), suggesting that the tail-length changes observed for cytoplasmic mRNAs were a consequence of limiting PABPC.

**Figure 4.**
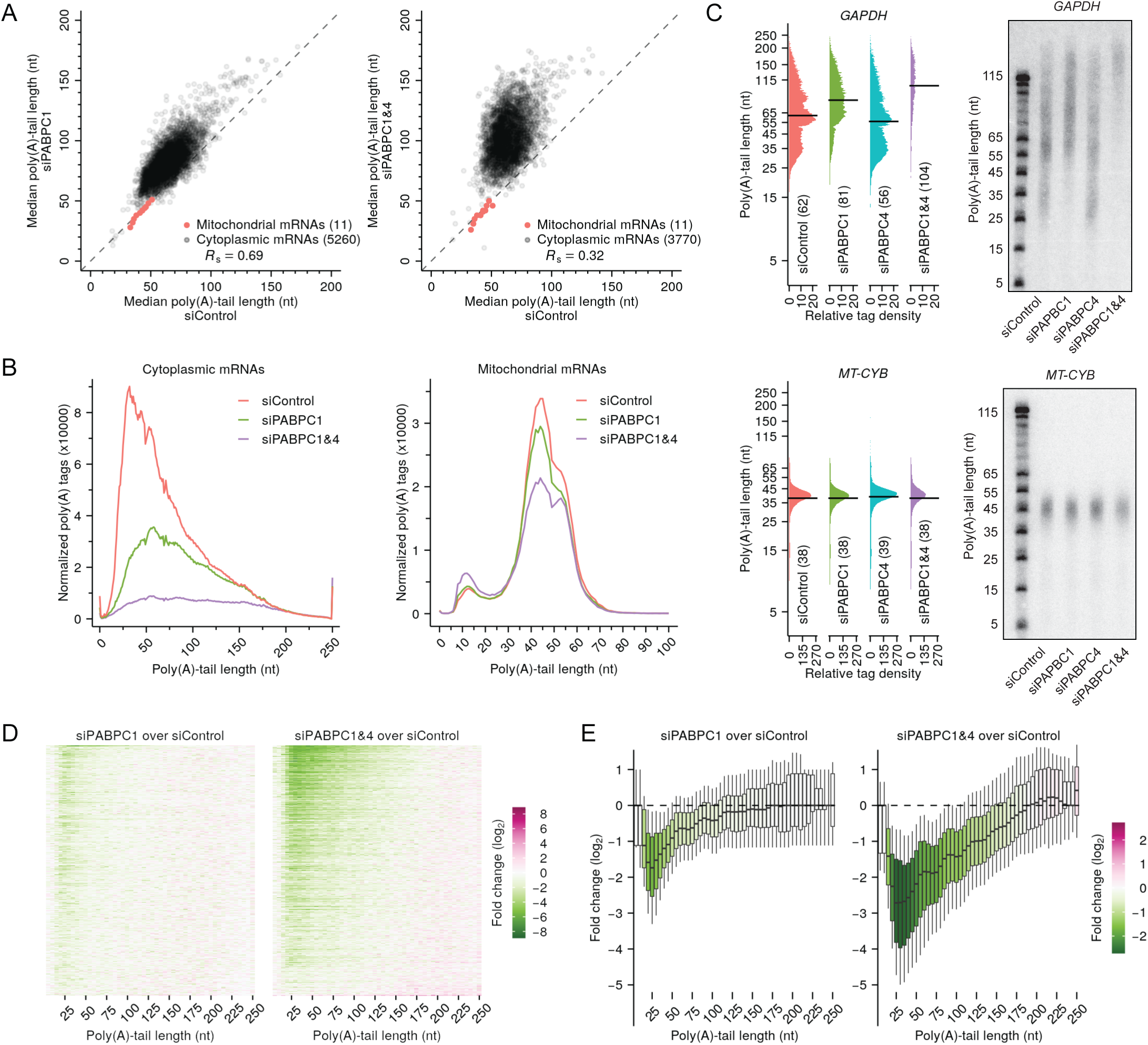
PABPC depletion causes premature decay of shorter-tail cytoplasmic mRNAs in HeLa cells. (**A**) The effect of PABPC knockdown on poly(A)-tail length. The plots compare median poly(A)-tail lengths in either PABPC1-knockdown cells (left) or PABPC1 and PABPC4 double-knockdown cells (right) to those in control cells. Results are shown for cytoplasmic mRNAs with ≥ 100 poly(A) tags (gray) and for mitochondrial mRNAs (red), merging data for *MT-ATP6* and *MT-APT8*, and for *MT-ND4* and *MT-ND4L*, which are bicistronic mitochondrial mRNAs. (**B**) The effect of PABPC knockdown on the abundance of mRNAs with different tail lengths. Shown are tail-length distributions of all cytoplasmic (left) and mitochondrial (right) mRNA poly(A) tags in control, PABPC1-knockdown, and double-knockdown cells. For each distribution, the abundance of tags was normalized to that of the spike-in tail-length standards. Due to depletion of tail-length calling at position 50, which was associated with a change in laser intensity at the next sequencing cycle, the values at this tail length were replaced with the average of values at tail lengths 49 and 51 nt. (**C**) The effect of PABPC knockdown on the abundance of mRNAs with different tail lengths, comparing of tail-lengths measured by sequencing (left) with those observed on RNase-H northern blots (right). Results are shown for a cytoplasmic mRNA *GAPDH* (top) and a mitochondrial mRNA *MT-CYB* (bottom). Relative tag density was calculated by log-transforming linear tag density using normalized poly(A) tag counts. Median tail-length values are indicated (horizontal lines) and listed in parentheses. For RNase-H northern blots, a DNA oligonucleotide complimentary to the 3′-UTR was used to direct cleavage of the target mRNA by RNase H, leaving a 35-nt fragment of the 3′-UTR appended to the poly(A) tail, which was resolved on a denaturing gel and detected by a radiolabeled probe. Tail lengths indicated along the left side of each gel are inferred from lengths of size markers. (**D**) The effect of PABPC knockdown on the abundance of mRNAs with different tail lengths, extending the intragenic analysis to tail-length distributions from thousands of genes. Heat maps compare poly(A)-tag levels in PABPC1-knockdown (left) or double-knockdown (right) cells to those in control cells, after normalizing to spike-in tail-length standards, as measured using tail-length sequencing. Each row represents mRNAs from a different gene, and rows are sorted based on fold change of mRNA abundance measured using RNA-seq. Only genes with ≥100 poly(A) tags in each of two samples being compared were included in the analyses (n = 5504). Columns represent values from 5-nt tail-length bins ranging from 0–244 nt and a 6-nt bin ranging from 245–250 nt. Tile color indicates the fold change of normalized tag counts (key). (**E**) The effect of PABPC knockdown on the abundance of mRNAs with different tail lengths, reanalyzing data from (**D**) to show distributions of poly(A)-tag changes observed at different tail-lengths. Each box-whisker shows the 10^th^, 25^th^, 50^th^, 75^th^, and 90^th^ percentile of fold changes in normalized poly(A)-tag counts observed for each tail-length bin of (**D**). The color of each box indicates the median value (key). The following figure supplements are available for figure 4: **Figure supplement 4**. PABPC depletion is not sufficient to establish strong coupling between poly(A)-tail length and TE in HeLa cells. **Figure supplement 5**. Support and extension of experiments showing that PABPC depletion causes premature decay of short-tailed mRNAs in HeLa cells.

We also used northern blots to examine effects of PABPC depletion on tail-length distributions of mRNAs from individual genes and found that the results corresponded well to those observed by tail-length sequencing. Both northern blots and sequencing showed strong depletion of short-tailed isoforms for a cytoplasmic mRNA (*GAPDH*) after PABPC knockdown but no substantial change in the tail-length distribution of a mitochondrial mRNA (*MT-CYB*) (Figure 4C). Using sequencing data to examine the intragenic tail-length distributions of cytoplasmic mRNA from each of more than five thousand other genes revealed findings resembling those observed for *GAPDH* (Figure 4D–E). After knocking down PABPC1, the reduction in short-tailed mRNAs was typically most severe for mRNA isoforms with tail lengths of ∼25 nt, and in the double knockdown, this dip at ∼25 nt became more pronounced, with reductions extending to all but the longest-tail isoforms (Figure 4D–E). Indeed, for more than half of the genes examined, ≥ 2-fold reductions extended to isoforms with tails as long as 135 nt (Figure 4D–E). Similar results were observed when examining tail-length distributions in the NIH3T3 dataset (Figure 4—figure supplement 5B), and when examining individual tail-length distributions for mRNAs of each of the top-expressed nuclear genes but not when examining those for mRNAs of mitochondrial genes (Figure 4—figure supplement 5F–G).

These results, which showed that mRNAs with short tails were preferentially destabilized when PABPC was depleted provided genetic loss-of-function evidence that PABPC stabilizes mRNAs in mammalian cell lines. Although genetic evidence for the role of PABPC in mRNA stability has been reported in yeast (Caponigro and Parker, 1995; Coller et al., 1998), this function had not been established in mammalian cells. Although in principle the mRNA destabilization that we observed upon PABPC-depletion might have been indirect, two lines of evidence support the conclusion that this destabilization was a direct consequence of the loss of PABPC binding to poly(A) tails. First, destabilization preferentially occurred for short-tailed mRNAs, which were expected to be the least successful at competing for binding under conditions of limiting PABPC. Second, destabilization sharply diminished at tail lengths of 10– 15 nt, which corresponded to the 12 nt poly(A) sequence reported to be the minimal length that can be bound by PABPC1 with high affinity (Kühn and Pieler, 1996). Indeed, the modest loss observed for mRNAs with very short poly(A) tails (Figure 4D–E) suggested that even in control cells that had abundant PABPC, mRNAs with tails shorter than 12 nt were poorly bound by PABPCs, and thus PABPC depletion did not substantially influence their abundance.

A recent study observed similar poly(A) tail-length changes in PABPC1-depleted cells but attributed these changes to impaired deadenylation (Yi et al., 2018). Because PABPC can promote deadenylation in vitro (Uchida et al., 2004; Webster et al., 2018; Yi et al., 2018; Schäfer et al., 2019), it is conceivable that the loss of PABPC would slow deadenylation, thereby increasing mRNA median tail lengths, as observed in Pab1-knockout yeast (Caponigro and Parker, 1995). However, our analyses, which had the benefit of quantitative tail standards that enabled measurement of absolute abundance changes, revealed little added accumulation of long-tailed isoforms in PABPC-depleted cells (Figures 4B–E, Figure 4—figure supplement 5B, 5F), indicating that a deadenylation defect was not the major cause for the perturbed tail-length distributions. Moreover, we found that mRNA half-life values, as determined by metabolic labeling, reduced significantly when PABPC was knocked down (Figure 5—figure supplement 6A–C), which concurred with the conclusion that PABPC knockdown destabilized short-tailed mRNA isoforms and argued against the previous assertion that PABPC knockdown impaired deadenylation, in that impaired deadenylation would have lengthened mRNA half-lives.

Taken together, these results show that PABPC binding stabilizes mRNAs of cultured mammalian cells; if PABPC becomes limiting in these cells, the short-tailed mRNAs become destabilized, presumably because they compete less effectively for PABPC. Most importantly, the destabilization of mRNAs that competed poorly for PABPC binding helps explain why limiting PABPC was insufficient to cause strong coupling in mammalian cell lines, in that strong coupling between tail length and TE would be difficult to establish in a regulatory regime in which short-tailed mRNA molecules that lack PABPC binding are degraded rather than translated less efficiently. Thus, these results identify a second mechanistic requirement for strong coupling between tail length and TE: In addition to limiting PABPC, strong coupling requires metabolic stability of the mRNAs that compete poorly for PABPC binding.

### Terminal uridylation contributes to premature decay of short-tailed mRNAs in PABPC-depleted cells

The identification of this second requirement for strong coupling brought to the fore the question of why mRNAs that competed poorly for limiting PABPC were destabilized. To explore the possible mechanisms, we searched for perturbations that could restore stability of short-tailed mRNAs in HeLa cells undergoing PABPC knockdown, monitoring tail-length distributions of endogenous *GAPDH* using northern blots. As a positive control, expressing an siRNA-resistant PABPC1 restored stability of short-tailed species, as did frog ePAB, which further illustrated functional conservation of PABPC from different species and developmental stages (Figure 5A). Interestingly, a PABPC1 variant with substitutions that disrupt its interaction with eIF4G (Chorghade et al., 2017) also restored the stability of short-tailed species, implying that the classical closed loop is not necessary for PABPC1 to protect short-tailed mRNAs from degradation. Because PABPC has been implicated in inhibiting mRNA terminal uridylation in vitro (Lim et al., 2014) and in cells (Yi et al., 2018), and terminal uridylation has been linked to mRNA decay (Rissland and Norbury, 2009; Lim et al., 2014), we asked if terminal uridylation contributed to the loss of short-tailed mRNAs. Knocking down both TUT4 and TUT7 in PABPC-depleted cells partially restored short-tailed *GAPDH* mRNAs (Figure 5B). Similar results were observed in HCT116 cells, in which we tagged endogenous PABPC1 with an auxin-inducible degron (AID) and induced depletion by adding indole-3-acetic acid (IAA, a form of auxin) (Figure 5C) (Natsume et al., 2016).

**Figure 5.**
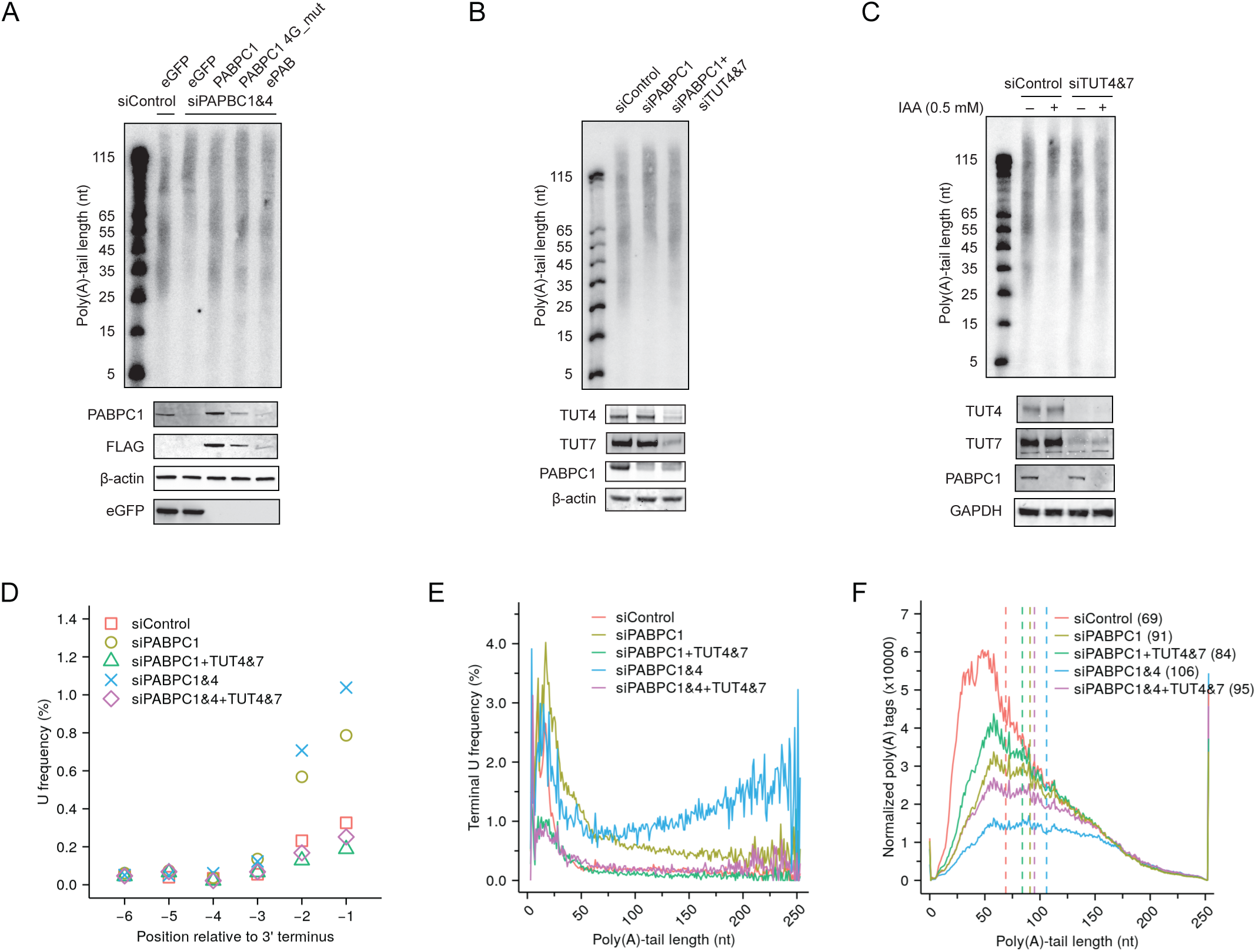
Depletion of TUT4 and TUT7 alleviates premature decay of mRNA caused by PABPC depletion. (**A**) Rescue of loss of shorter-tail mRNAs in PABPC-depleted HeLa cells by expressing either an siRNA-resistant human *PABPC1*, an siRNA-resistant *PABPC1* coding for a mutant that does not bind eIF4G (4G_mut), or an siRNA-resistant frog *ePAB*. At the top is an RNase H northern blot probed for *GAPDH*, as in Figure 4C. At the bottom is a western blot probed for the indicated proteins or the FLAG tag appended to the C terminus of each of the expressed PABPC proteins. (**B**) Partial rescue of loss of shorter-tail mRNAs in PABPC-depleted HeLa cells by knocking down *TUT4* and *TUT7*. At the top is an RNase H northern blot probed for *GAPDH*, as in Figure 4C. At the bottom is a western blot probed for the indicated proteins. (**C**) Partial rescue of loss of shorter-tail mRNAs in PABPC-depleted HCT116 cells by knocking down of *TUT4* and *TUT7*. Endogenous PABPC1 was tagged with AID, and IAA was added for 24 h to target the fusion protein for degradation. Otherwise, this panel is as in (**B**). (**D**) The effect of PABPC knockdown and TUT knockdown on terminal uridylation. Plotted is the fraction of uridines near the termini of cytoplasmic mRNAs in HeLa cells transfected with the indicated siRNAs, as measured using tail-length sequencing. (**E**) The effect of PABPC knockdown and TUT knockdown on terminal uridylation of tails with different lengths. Shown is terminal uridylation frequency of cytoplasmic mRNAs as a function of poly(A)-tail length in HeLa cells transfected with the indicated siRNAs, as measured using tail-length sequencing. (**F**) Partial rescue of loss of shorter-tail cytoplasmic mRNAs in PABPC-depleted HeLa cells by knocking down *TUT4* and *TUT7*, extending the analysis to all tail-length sequencing tags. Plotted are tail-length distributions from HeLa cells transfected with the indicated siRNAs. For each distribution, tag counts were normalized to spike-in tail-length standards. Median values are indicated (dashed lines) and listed in parentheses. The following figure supplement is available for figure 5: **Figure supplement 6**. Measurements of mRNA half-lives in PABPC-depleted HeLa cells and terminal uridylation levels in frog oocytes.

To examine the global rescue of short-tailed mRNAs and at the same time monitor mRNA terminal uridylation levels, we modified our tail-length sequencing protocol by including in the adaptor-ligation step a splint oligonucleotide designed to accommodate tails with a 3′ terminal U (Eisen et al., 2020). Knockdown of PABPC1 alone significantly increased the terminal uridylation levels across essentially all tail-length isoforms (Figure 5D–E), consistent with a previous report (Yi et al., 2018). Knockdown of PABPC4 in addition to PABPC1 further increased uridylation of mRNA isoforms with longer tail-lengths (Figure 5E). Knockdown of TUT4 and TUT7 in PABPC-depleted cells brought terminal uridylation of all tail-length isoforms to background levels (Figure 5D–E) and, more importantly, preferentially rescued shorter-tail isoforms, thereby decreasing median tail lengths (Figure 5F). These results were consistent with those of our northern assays, and together, our results indicated that in these mammalian cells, limiting PABPC makes short- and medium-tailed mRNA isoforms that poorly compete for PABPC binding more susceptible to terminal uridylation, thereby accelerating their decay.

### PABPC has little effect on TE in mammalian cell lines

Having found that destabilization of short-tailed mRNAs dampened coupling between poly(A)-tail length and TE in PABPC-depleted mammalian cells, a key question remained regarding how mRNA TE, if not influenced by poly(A)-tail length, was affected in these cells. To answer this question, we conducted global profiling of HeLa cells 48 h after siRNA transfection, a time point at which PABPC1 and PABPC4 knockdowns were substantial but secondary effects were presumably not yet too severe (Figure 6—figure supplement 7A). We also modified our RNA-seq and ribosome-profiling protocols to enable absolute TE comparison by using mitochondrial mRNAs for normalization (Rooijers et al., 2013). Surprisingly, near-complete depletion of PABPCs had no detectable effect on global mRNA TEs (Figure 6A–B). Although the protein synthesis rate in double-knockdown cells was reduced to 75.7% of that observed in control cells, as measured by averaging ribosome footprints changes observed for the 9,697 analyzed genes (Figure 6A–B), which agreed with results of a global puromycin-based translation assay (78.7%) (Figure 6C), this reduced protein synthesis was fully explained by the decrease in mRNA levels, as indicated by a distribution of TE changes that centered near zero (Figure 6A–B). Examination of our ribosome-footprinting data revealed no upregulation of other PABPC paralogs, although the TE of *PABPC1* mRNA increased 2.5-fold (Figure 6A), consistent with its known autoinhibitory translational control (Bag and Wu, 1996). Results of polysome profiling confirmed those of our sequencing-based methods, in that the reduction in translation output, as measured by the height of polysome peaks in double-knockdown cells, was attributable to an overall decrease of mRNA levels rather than to decreased TEs that would otherwise cause a shift of mRNA distribution from heavy to light fractions (Figure 6—figure supplement 7B). Together, these results indicated that PABPC depletion in HeLa cells had negligible effect on mRNA TE.

**Figure 6.**
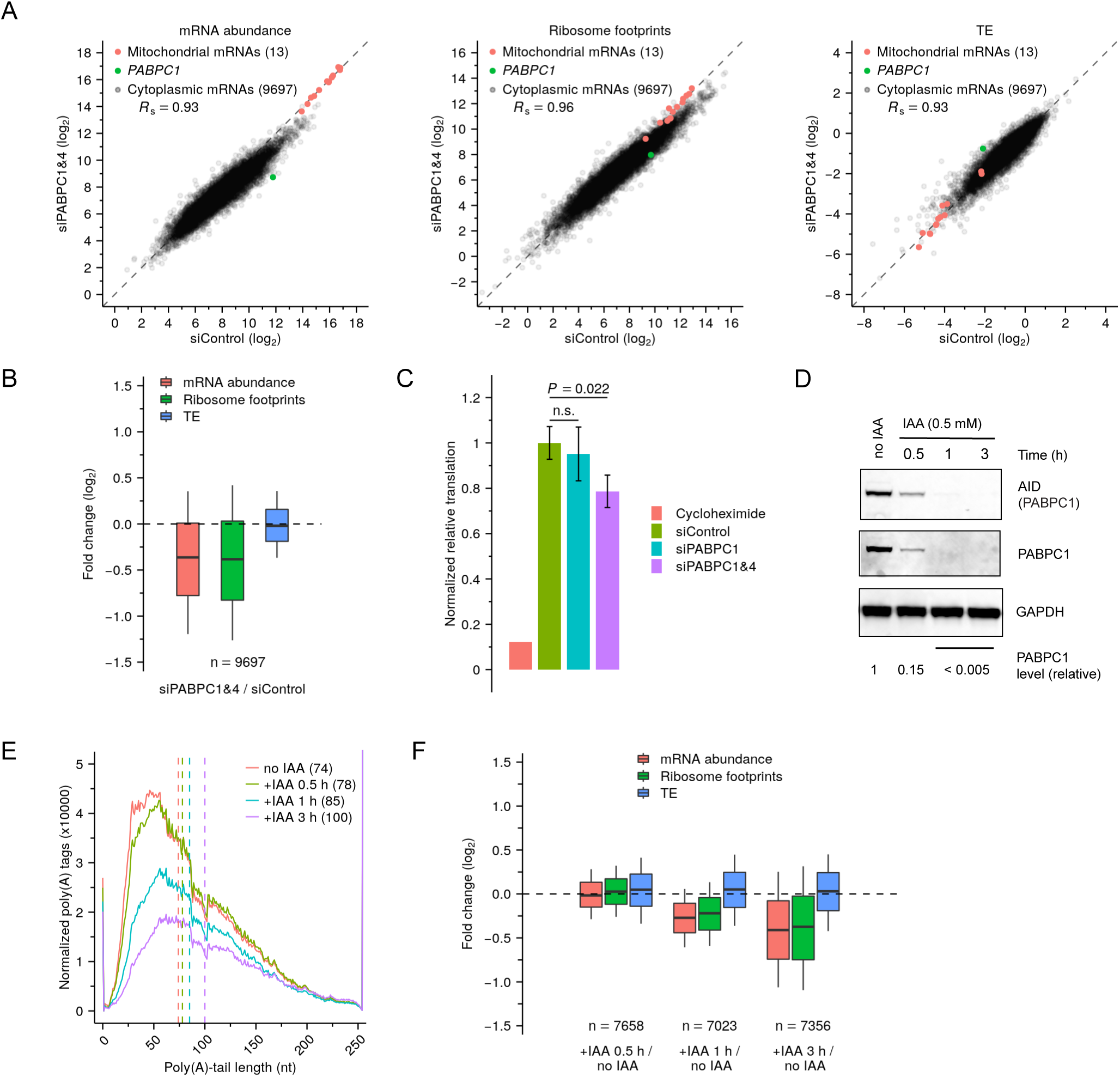
Depletion of PABPC in mammalian cell lines has minimal effect on TE. (**A**) Effect of PABPC knockdown on mRNA abundance (left), ribosome-footprint abundance (middle) and TE (right) in HeLa cells, comparing values in double-knockdown cells to those in control cells. For each gene, the number of mRNA and ribosome-footprint reads per kilobase are shown after normalizing to measurements for mitochondrial mRNAs. (**B**) Distributions of the effects of PABPC double knockdown on mRNA abundance, ribosome-footprint abundance and TE. Each box-whisker shows the 10^th^, 25^th^, 50^th^, 75^th^ and 90^th^ percentile of the fold changes observed in (**A**). (**C**) Effect of PABPC knockdown on protein synthesis in HeLa cells. Plotted are relative levels of protein synthesis measured by pulse puromycin incorporation 48 h after transfection with the indicated siRNAs (Figure 6—figure supplement 7A; error bars, standard deviation of three biological replicates; *P* values, one-sided *t*-tests; n.s., not significant). (**D**) Western blot showing rapid degradation of PABPC1-AID fusion protein after adding IAA to genome-engineered HCT116 cells. PABPC1 protein levels were quantified by averaging signals for AID and PABPC1, after normalizing to that for GAPDH. (**E**) Effect of PABPC1 depletion on abundance of mRNAs with different tail lengths. Shown are tail-length distributions of all poly(A) tags obtained from cytoplasmic mRNAs of HCT116 PABPC1-AID cells after treatment with IAA for the indicated time. For each distribution, the abundance of tags was normalized to that of the spike-in tail-length standards. Median values are indicated (dashed lines) and listed in parentheses. Due to depletion of 101-nt tail lengths, which was associated with a change in laser intensity at the next sequencing cycle, the values at this length were replaced with the average of values at tail lengths 100 and 102 nt. (**F**) Summary of the effects of rapid PABPC1 depletion. For each gene, at each time point after adding IAA to HCT116 PABPC1-AID cells, values for mRNA abundance, ribosome-footprint abundance and TE were each compared to the value observed in cells not treated with IAA (Figure 6—figure supplement 7C), and the distribution of fold changes is plotted. Each box-whisker shows the 10^th^, 25^th^, 50^th^, 75^th^ and 90^th^ percentile. The following figure supplement is available for figure 6: **Figure supplement 7**. Depletion of PABPC in mammalian cell lines has minimal effect on TE.

The slow dynamics of siRNA-mediated knockdown, which dictated the relatively late 48 h time point for our global measurements, might have prevented detection of any TE changes that happened earlier, before a new steady state had been reached. To examine this possibility, we monitored the dynamics of tail-length, mRNA-abundance, and translation changes soon after PABPC depletion, using the HCT116 PABPC1-AID degron cell line, in which PABPC1 was rapidly and efficiently depleted after adding IAA (85% within 30 min, > 99% within 1 h, Figure 6D). Because PABPC1 is the primary PABPC isoform in HCT116 cells, depletion of PABPC1 alone caused substantial destabilization of shorter-tail mRNAs 1 h after IAA addition (Figure 6E), and this destabilization further increased after 3 h. Accompanying the loss of shorter-tail mRNAs was a corresponding reduction of mRNA abundance for most genes (Figure 6F, Figure 6—figure supplement 7C). Importantly, ribosome footprints declined in lockstep with mRNA abundance, leading to median TE changes that centered near zero over the entire course of PABPC1 depletion (Figure 6F, Figure 6—figure supplement 7C). Thus, as PABPC became limiting, mRNAs that lost PABPC had no detectable reduction in TE before they were destabilized.

These results show that in contrast to mRNAs of frog oocytes and presumably those of other coupled systems, mRNAs of HeLa and HCT116 cells do not require PABPC for efficient translation, which explains why poly(A)-tail lengths were not able to strongly influence TE after we reduced PABPC of these cells to limiting levels. Thus, these results identify a third important mechanistic requirement for coupling between poly(A)-tail lengths and TE: coupling requires a regulatory regime in which PABPC affects mRNA translation.

## Discussion

We find that three fundamental molecular conditions must be met for cells to use poly(A)-tail lengths to effectively regulate TE. First, PABPC must be limiting compared to the number of poly(A) sites available for binding. Under this condition, short-tailed mRNA isoforms that poorly compete for PABPC are less likely to have PABPC bound to their 3′ ends (Figure 7). To the extent that PABPC is not bound, these isoforms lose the translation-activating capability of PABPC observed in coupled systems. When additional PABPC is introduced into a coupled system, short-tailed mRNAs benefit more from PABPC binding than long-tailed ones, which are more likely to already possess the number of PABPC molecules required for more efficient translation. At saturating PABPC, both short and long tails are presumably coated by PABPC (Figure 7). Although long-tailed mRNAs will have more PABPC molecules bound, the ability to promote translation might be conferred by a single bound PABPC, with much less added benefit conferred by additional binding. In this scenario, the strong influence of poly(A)-tail lengths on TE disappears.

**Figure 7.**
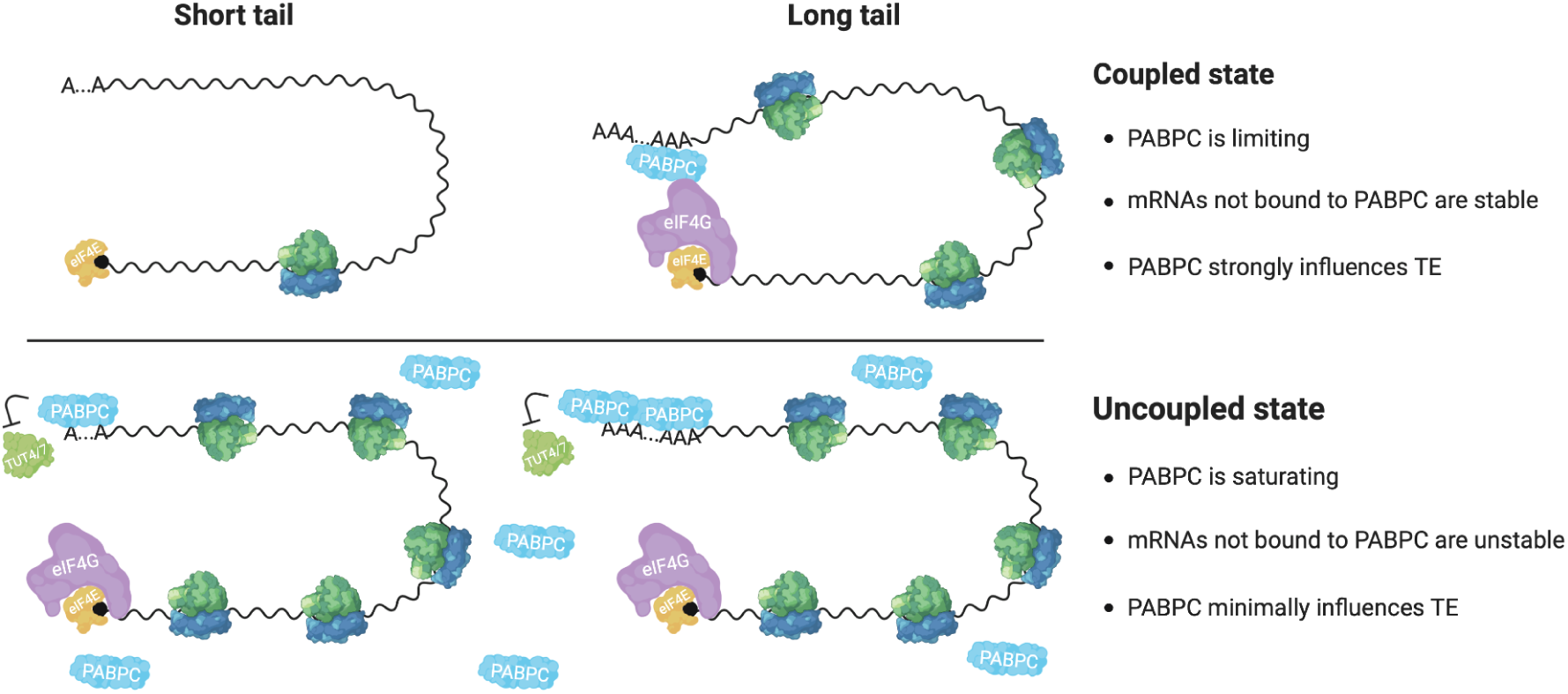
Model for coupling between poly(A)-tail length and TE, and context-dependent roles of PABPC. See text for details.

The requirement of limiting PABPC for coupling TE to poly(A)-tail length raises the question of the stoichiometry between PABPC and its sites in the poly(A) tails and whether this stoichiometry changes during the embryonic switch of gene-regulatory regimes. Our sequencing results indicated that the mRNAs of a stage VI frog oocyte have ∼2.8 x 10^11^ PABPC sites (∼7 x 10^10^ sites per 1 µg total RNA, ∼4 µg total RNA per oocyte), which concurred with the previous estimate of 2∼3 x 10^11^ sites per oocyte (Sagata et al., 1980). The amount of PABPC (including both ePAB and PABPC1) in frog oocytes has been estimated at either 1 x 10^11^ (Cosson et al., 2002) or 1.4 x 10^11^ (Smits et al., 2014; Peuchen et al., 2017) molecules per oocyte. Our results showing that PABPC activity is limiting in frog oocytes, agreed with these estimates that imply that PABPC levels are sufficient to bind no more than half of the available PABPC sites. Poly(A)-site occupancy would be even lower if some PABPC proteins were sequestered from the mRNA pool or in an inactive form, which could be conferred by post-translational modifications that affect the ability of PABPC to either bind poly(A) tails or promote translation (Brook et al., 2012). In our overexpression experiment, we increased the level of ePAB to twice that of its endogenous level (Figure 3—figure supplement 3B). Although this increase was substantial, the overall PABPC level was unlikely to saturate all poly(A) sites, which explains why the coupling between poly(A)-tail length and TE diminished but was not completely lost (Figure 2A, 3C–E). Until stage 15, developing frog embryos maintain the number of PABPC sites at a level resembling that of oocytes (Sagata et al., 1980). In contrast, the total amount of PABPC molecules increases significantly, nearly tripling by stage 12 (Peshkin et al., 2019)—the stage at which the coupling between tail length and TE starts to disappear (Subtelny et al., 2014). This increased PABPC would shift the stoichiometry towards PABPC being less limiting. Importantly, dysregulation of this tightly controlled stoichiometry not only disrupts the normal gene-regulatory regime but also can cause severe consequences during oocyte maturation and embryonic development (Wormington et al., 1996; Gorgoni et al., 2011).

Although translation of many short-tailed mRNAs improved substantially when PABPC was overexpressed in frog oocytes, translation of some failed to improve. Perhaps these mRNAs were sequestered away from the translation machinery in the cytoplasm and thus unable to benefit from overexpressed PABPC proteins. One likely location for such sequestration would be germ granules, which have been implicated in regulating mRNA translation in oocytes of diverse animal species (Voronina et al., 2011). This mechanism might reinforce coupling between poly(A)-tail length and TE, perhaps by selectively sequestering short-tailed mRNAs from the active translation pool.

The second condition required for strong coupling between poly(A)-tail length and TE is the survival of short-tailed mRNAs under conditions in which PABPC is limiting (Figure 7). This condition was not met in the post-embryonic mammalian cell lines we examined. When PABPC was depleted in these uncoupled systems, many mRNA molecules, particularly short-tailed ones that presumably competed poorly for the remaining PABPC, were degraded. The preferential loss of mRNAs not bound by PABPC reduced the range of tail lengths, which correspondingly reduced the range of TEs that could potentially be imparted by coupling between tail length and TE. This reduced range also presumably reduced the ability to detect coupling, although some ability was expected to be retained, as indicated by an analysis in which data from a coupled system was sampled to match the more restricted tail-length distribution of an uncoupled system (Subtelny et al., 2014). More importantly, the loss of mRNAs not bound by PABPC reduced the number of PABPC-binding sites, thereby reducing the extent to which these sites were in excess over PABPC and thus reducing coupling between tail length and TE.

In oocytes and early embryos, mRNA decapping is uncoupled from deadenylation (Gillian-Daniel et al., 1998), which helps explain why short-tailed mRNAs survive in these systems despite our finding that they have limiting PABPC activity. We suggest two nonexclusive mechanistic explanations for this unusual decoupling of decapping from deadenylation. First, oocytes and early embryos have relatively low expression of decapping enzymes (Ma et al., 2013; Peshkin et al., 2019). Second, mRNA terminal uridylation activity is very low in frog oocytes (Figure 5—figure supplement 6D) (Chang et al., 2018), and it remains low throughout early embryonic development and only starts to increase dramatically after zygotic genome activation (Chang et al., 2018). Because terminal uridylation plays an important role in degrading short-tailed mRNAs that lack PABPC binding in mammalian cells (Figure 5) (Lim et al., 2014; Yi et al., 2018), this developmental delay of terminal uridylation might help ensure the survival of short-tailed mRNAs and thereby enable strong coupling between poly(A)-tail length and TE in oocytes and early embryos.

The third condition required for coupling between tail length and TE is that PABPC must have the ability to influence TE of bound mRNAs (Figure 7). In the coupled system of frog oocytes, increasing PABPC levels substantially improved TE of nearly all mRNAs (Figure 2B–E). In contrast, in uncoupled systems such as HeLa and HCT116 cells, severe depletion of PABPC, such that short- and medium-tailed mRNAs were markedly destabilized, had no consistent impact on mRNA TE (Figure 6B and F). We suspect that the differential effect of PABPC on TE observed in coupled and uncoupled systems is related to the divergent levels of basal translation initiation observed between these systems. Indeed, the overall translation measured by polysome profiles in oocytes and early embryos is much lower than that observed in either later developmental stages (Woodland, 1974) or post-embryonic mammalian cell lines (Figure 3—figure supplement 3A, Figure 6—figure supplement 7B), which provides the opportunity for a translation-activating effect of PABPC to be more prominent in the coupled systems.

Our results showing that PABPCs, while playing a crucial role in protecting mRNA from premature decay, have minimal contribution to translation in post-embryonic mammalian cell lines might seem to contradict the well-accepted function of PABPC as a translational activator. However, many previous studies that established the role for PABPC in promoting translation were carried out in frog oocytes or early embryonic systems (Smith et al., 2014), where we found PABPC to globally enhance TE. Other previous studies were conducted in vitro, with mixed results: in reconstituted systems, PABPC is dispensable for translation initiation (Mitchell et al., 2010), and in rabbit reticulocyte lysates, PABPC has a minimal effect on translation (Hinton et al., 2007), whereas in some other cell extracts, PABPC activates translation (Tarun and Sachs, 1995; Kahvejian et al., 2005). Additional experiments will be required to determine whether this discrepancy between results we obtained from living cells and those previously obtained in some post-embryonic cell extracts are attributable to differences between cell types or to differences between cellular cytoplasm and in vitro extracts—perhaps imparted by dilution of translational components in extracts. Such studies will need to differentiate between the translation-activation and mRNA-stabilization activities of PABPC. In the meantime, it is helpful to know that PABPC stabilizes mRNAs of post-embryonic metazoan cells and that the two activities of PABPC can be context dependent, such that in frog oocytes PABPC strongly activates translation and has no effect on mRNA stability, whereas in mammalian cell lines it stabilizes mRNAs and has no detectable effect on TE.

The dual potential of PABPC in stabilizing mRNA and promoting translation bestows PABPC with distinct and context-dependent roles in regulating protein synthesis. In oocytes and early embryos, the lack of mRNA transcription and degradation leaves differential TE as the primary option for modulating protein synthesis. In this context, limiting PABPC proteins bind primarily to mRNAs with longer poly(A) tails and activate their translation. In contrast, in post-embryonic mammalian cell lines, where transcription, mRNA degradation, and translation are each operating at high efficiency, PABPCs protect mRNAs from pre-mature decay, enabling them to contribute the proper amount of protein during their lifetimes, but without any additional enhancement of TE. These context-dependent activities are not only crucial for understanding how coupling between tail length and TE is established in oocytes and early embryos and why it is lost later in development, they also provide mechanistic insight into the effects of the many posttranscriptional regulatory phenomena that alter poly(A)-tail lengths. For example, during miRNA-mediated repression, the Argonaute–miRNA complex binds target mRNAs and recruits factors that displace PABPC from the poly(A) tail and accelerate tail shortening (Giraldez et al., 2005; Wu et al., 2006; Fabian et al., 2009; Moretti et al., 2012; Rissland et al., 2017). In early zebrafish embryos, these effects are expected to disadvantage target mRNAs at competing for limited PABPC, which in this context would reduce their TE without changing their stability, thereby explaining why miRNAs primarily cause translational repression in these early embryos (Bazzini et al., 2012; Subtelny et al., 2014). However, in later embryonic development as well as in post-embryonic mammalian cells, displacement of PABPC and accelerated tail shortening would reduce PABPC binding to target mRNAs, which in this context would reduce their stability without changing their TE, thereby explaining why miRNAs primarily cause mRNA destabilization in these cells (Guo et al., 2010; Bazzini et al., 2012; Subtelny et al., 2014).

How PABPC promotes translation remains an enigma, although the closed-loop model has offered a sound mechanistic explanation. The interaction between eIF4G and PABPC is well characterized and provides a physical link connecting both ends of the mRNA, and some studies are able to catch a glimpse of possible circular structures of mRNAs in fixed tissues (Christensen et al., 1987) or in vitro (Wells et al., 1998). However, a recent single-molecule imaging study in mammalian cells questions the widespread existence of mRNA closed-loop structures (Adivarahan et al., 2018). Our finding that depletion of PABPC had minimal impact on TE in mammalian cell lines supports a model in which pervasive eIF4G–PABPC-associated looping of mRNAs is generally lacking in uncoupled systems. This idea is consistent with the finding that the interaction between eIF4G and PABPC is dispensable in yeast (Park et al., 2011) and HEK293 cells (Adivarahan et al., 2018), both of which are uncoupled systems (Subtelny et al., 2014). In contrast, the eIF4G–PABPC interaction is critical during frog oocyte maturation (Wakiyama et al., 2000), a process that relies on coupling between poly(A)-tail length and TE (Richter and Lasko, 2011), and our results showed that increasing PABPC had a global effect on upregulating TE in frog oocytes. Thus, the closed-loop structure, if it exists, might be more prevalent in coupled systems, such as oocytes or early embryos.

Our experiments focused on cells that inherently possessed either all or none of the three molecular conditions that we found to be required for strong coupling between tail length and TE, i.e., limiting PABPC activity, stabilization of mRNAs lacking a bound PABPC, and PABPC-sensitive translation. Nonetheless, some cells might fall in between these two extremes, processing one or two of these conditions but not all three. The results of our experiments in which we removed one of the conditions from frog oocytes or imposed one or two of the conditions in post-embryonic mammalian cells, predict that such cells that inherently fall between the two extremes have minimal if any coupling. Indeed, the concept that multiple conditions must be met before strong coupling can be established helps to explain why coupling between tail length and TE has been so infrequently detected outside the gene regulatory regime operating in oocytes and early embryos (Subtelny et al., 2014).

## Acknowledgments

We thank S. Eichhorn, T. Eisen, A. Subtelny, S. Gupta, X. Wu, S. McGeary, W. Fang, K. Lin, J. Smith, J. Kwasnieski and H. Sive for valuable discussions; S. Eichhorn, T. Eisen, K. Lin, and S. Gupta for sharing improved methods for poly(A)-tail profiling; K. McKinley, K. Su, M. Kanemaki, A. Holland, A.-B. Shyu for sharing cell lines and plasmids; N. Gray and M. Brook for antibodies, and the Whitehead Institute Genome Technology Core for sequencing. This work is supported by NIH grant GM118135. K.X. is a Cancer Research Institute Irvington Fellow supported by the Cancer Research Institute. D.P.B. is an investigator of the Howard Hughes Medical Institute.

## Author contributions

K.X. and D.P.B. conceived the project, designed the study, and wrote the manuscript. K.X. performed the experiments and the associated analyses.

## Figure legends

**Figure supplement 1.**
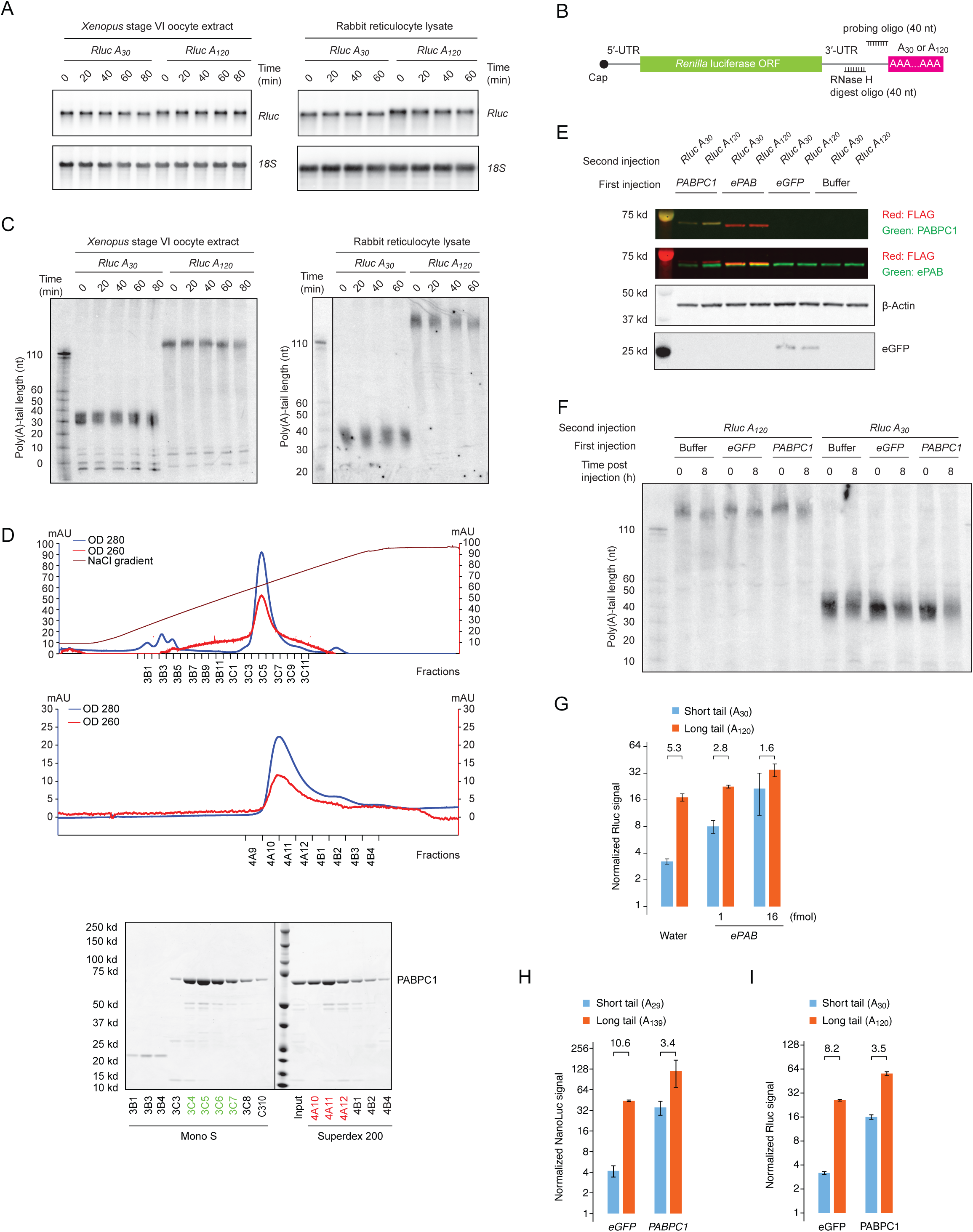
Supporting data for reporter experiments examining the effect of PAPBC levels on coupling between tail length and translation. (**A**) The stability of reporters during in vitro translation. Shown are mRNA northern blots monitoring reporter mRNA levels over the course of the indicated in vitro translation reactions. (**B**) Schematic of mRNA cleavage and probing strategy used in RNase H northern blots, which were used to examine tail lengths. A DNA oligonucleotide complimentary to the 3′-UTR was used to direct cleavage of the target mRNA by RNase H, leaving a 40-nt fragment of the 3′-UTR appended to the poly(A) tail, which was resolved on a denaturing gel and detected by a radiolabeled probe. (**C**) The integrity of reporter poly(A) tails during in vitro translation. Shown are RNase H northern blots monitoring poly(A)-tail lengths of reporter mRNAs in in vitro translation reactions, as illustrated in (**B**). Reference tail lengths are inferred from lengths of size markers. (**D**) Purification of PABPC1. At the top is a chromatogram of cation exchange after nickel affinity columns. In the middle is a chromatogram of gel filtration after the cation exchange. At the bottom are Coomassie-stained SDS-PAGE gels monitoring fractions collected during the cation exchange (Mono S) and the gel filtration (Superdex 200). Green fractions were pooled after the cation-exchange step, and red fractions were pooled after the gel-filtration step. (**E**) Expression of proteins from mRNAs injected into frog oocytes. Shown is a western blot detecting proteins from frog oocytes in which mRNAs were injected for expressing the indicated proteins. (**F**) The integrity of reporter tail lengths after injection into oocytes. Shown is an RNase H northern blot monitoring poly(A)-tail lengths of reporter mRNAs injected into oocytes. Otherwise this panel is as in (**C**). (**G**) The preferential effect of PAPBC1 and ePAB overexpression on translation of the short-tailed *Rluc* reporter in frog oocytes. Shown are relative luciferase signals following serial injection of mRNAs into frog oocytes, as in Figure 1D. Differential expression of PABPC1 or ePAB was achieved by injecting the indicated amount of mRNA in the first injection. The number above each bracket indicates the fold difference (error bars, standard deviations from three biological replicates). (**H**) The preferential effect of PABPC overexpression on translation of the short-tailed *NanoLuc* reporter in frog oocytes. Shown are relative luciferase signals from *NanoLuc* reporter mRNAs in frog oocytes overexpressing eGFP or PABPC1 in a serial-injection assay, as in Figure 1D. Otherwise, this panel is as in (**G**). (**I**) The preferential effect of PABPC protein injection on translation of the *Rluc* short-tailed reporter in frog oocytes. Shown are relative luciferase signals of *Rluc* reporter mRNAs co-injected with recombinant eGFP or PABPC1 proteins in frog oocytes. Otherwise, this panel is as in (**G**).

**Figure supplement 2.**
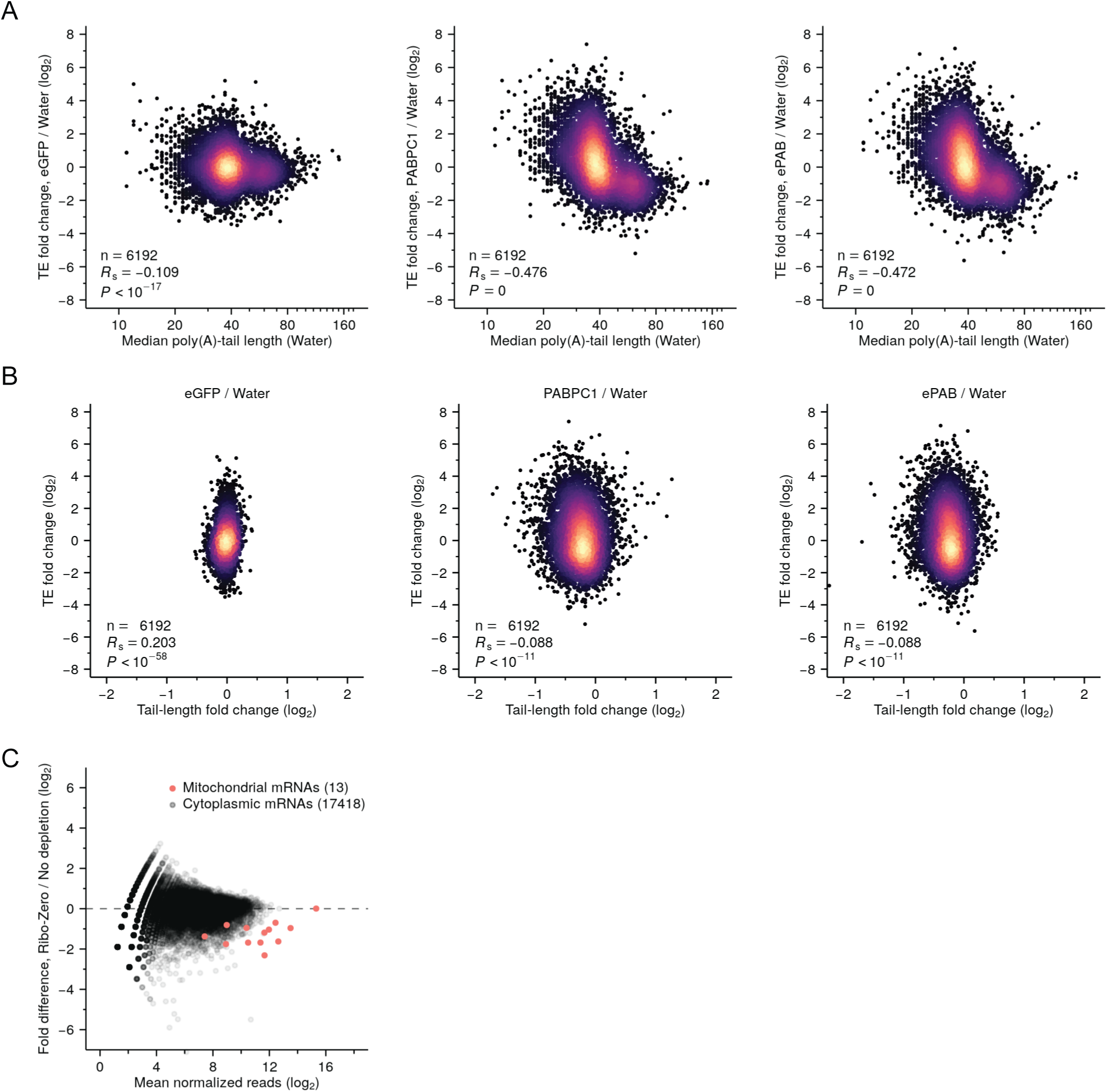
Increased PABPC increases translation of endogenous short-tailed mRNAs in frog oocytes. (**A**) Increased relative translation of short-tailed mRNAs after overexpressing PABPC in oocytes. Shown for each gene with ≥ 100 poly(A) tags is the change in relative TE observed after overexpressing either eGFP, PABPC1 or ePAB, plotted as a function of median poly(A)-tail length in water-injected oocytes. (**B**) Poor correspondence between TE changes and tail-length changes after overexpressing PABPC in oocytes. TE changes of (A) are plotted as a function of median poly(A) tail-length changes. (**C**) Depletion of frog mitochondrial mRNAs by the Illumina Ribo-Zero Gold kit.

**Figure supplement 3.**
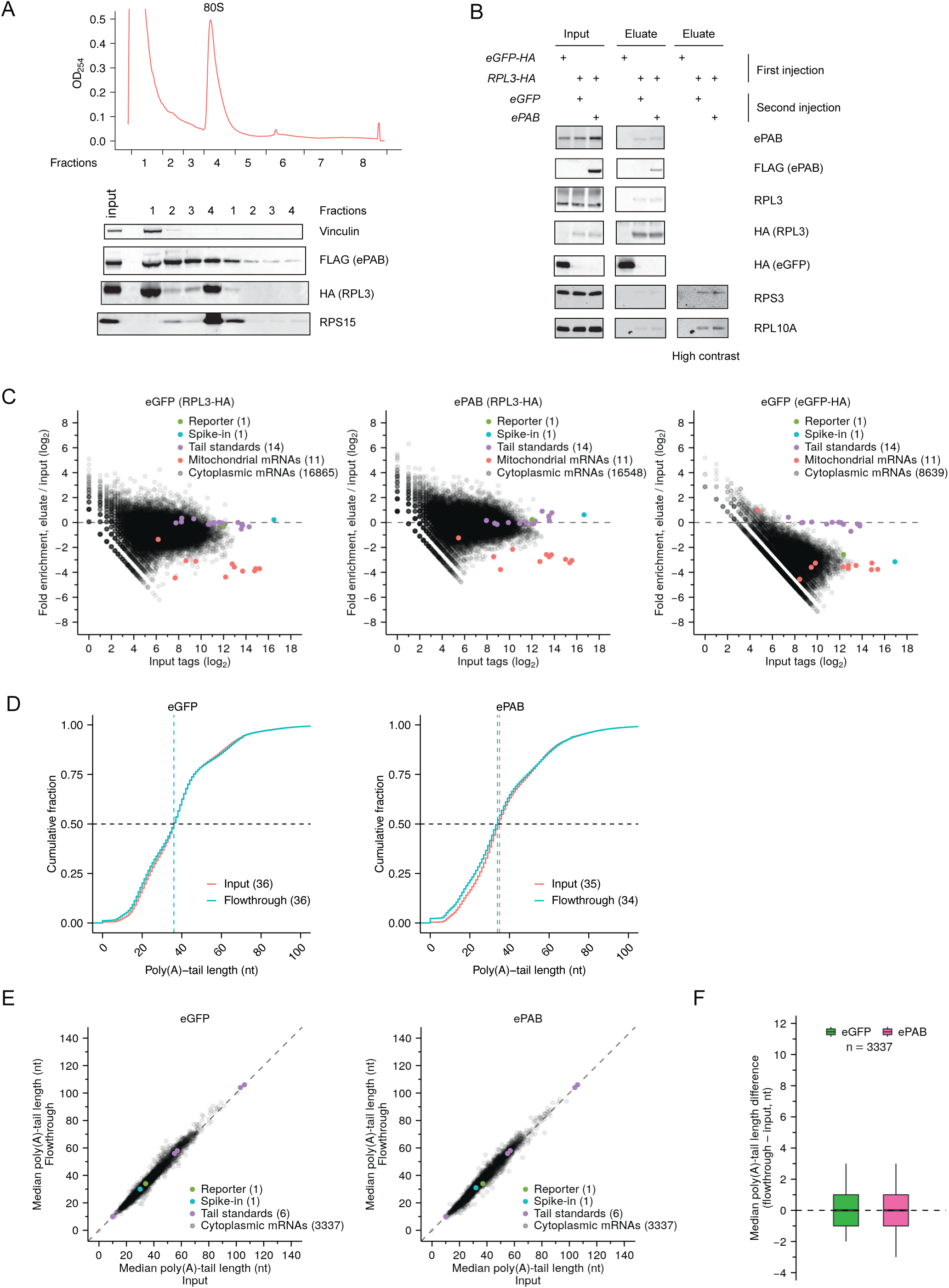
Supporting data for PAL-TRAP analyses in frog oocytes. (**A**) Incorporation of HA-tagged RPL3 into ribosomes. At the top is a sucrose-gradient polysome profile from frog oocytes injected with an mRNA encoding HA-tagged RPL3. At the bottom is a western blot of fractions collected from the polysome profile and probed for the indicated proteins. (**B**) Western-blot analysis of proteins in the input and eluate of PAL-TRAP experiments. Labels at the top show mRNAs that were injected into oocytes to overexpress the indicated proteins. (**C**) Fold enrichment in the PAL-TRAP analysis (eluate over input) as a function of expression (tags in the input) for each mRNA isoform with a unique 3′ end. Poly(A) tags were normalized to the spike-in tail-length standards. Poly(A) tags for bicistronic mitochondrial mRNAs (*mt-atp6* and *mt-apt8*, and *mt-nd4* and *mt-nd4l*) were merged. Also shown are results from a co-injected reporter mRNA, a spike-in mRNA and tail-length standards, as in Figure 3D. Poly(A) tags were normalized to the spike-in tail-length standards. Cytoplasmic mRNAs were enriched by HA-IP in RPL3-HA-expressing oocytes, as shown by relative depletion of mitochondrial mRNAs, which are not translated by cytoplasmic ribosomes. In contrast, almost all cytoplasmic mRNAs were depleted by HA-IP in eGFP-HA-expressing oocytes. (**D**) The similarity of PAL-TRAP results for input and flowthrough when pooling data for all mRNA isoforms. Otherwise, this panel is as in Figure 3C. (**E**) The similarity of PAL-TRAP results for input and flowthrough when examining data for individual mRNA isoforms. Otherwise, this panel is as in Figure 3D. (**F**) Summary of differences in median tail lengths observed between the flowthrough and the input for mRNA isoforms represented in (**E**). Box and whiskers indicate the 10^th^, 25^th^, 50^th^, 75^th^ and 90^th^ percentiles.

**Figure supplement 4.**
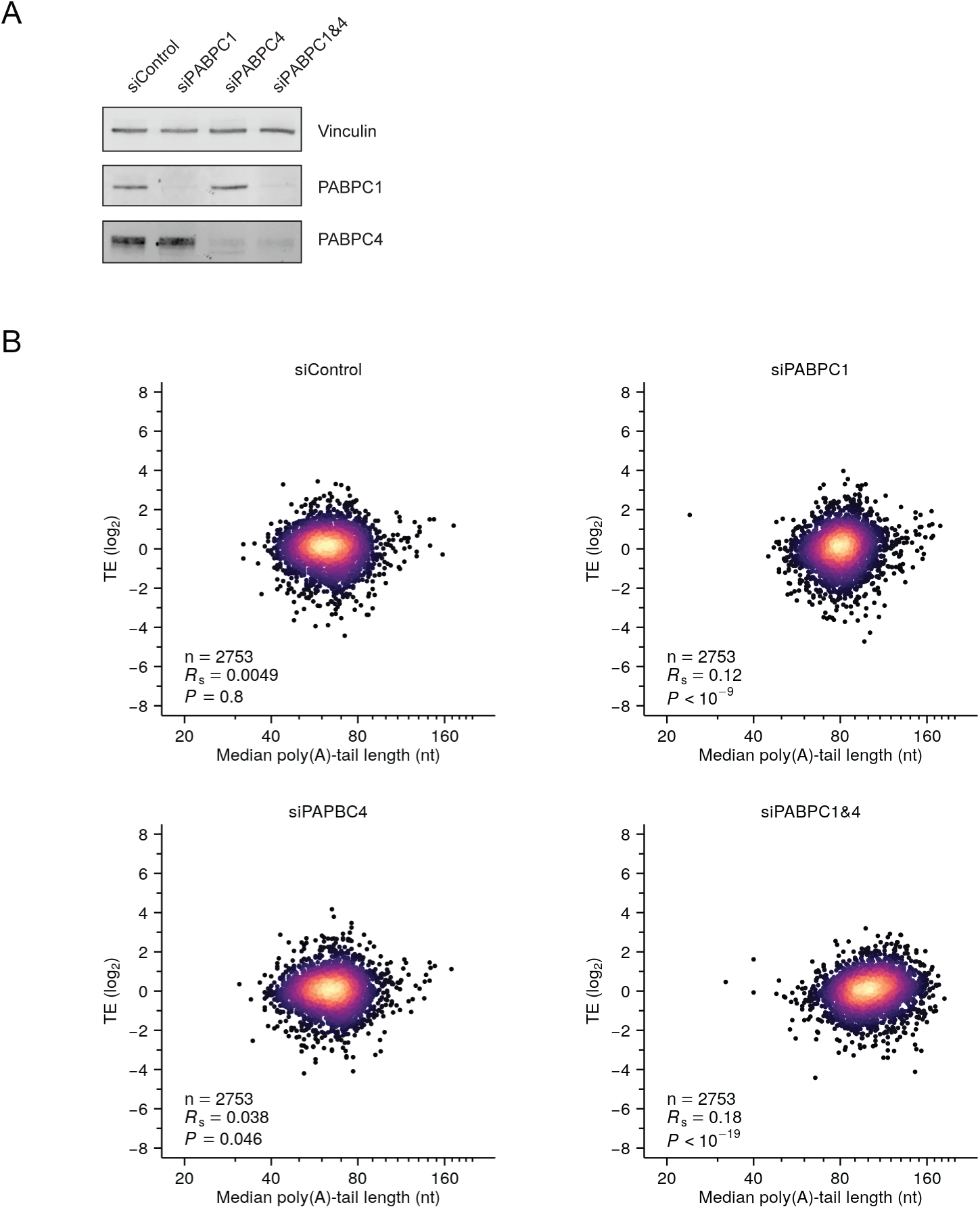
PABPC depletion is not sufficient to establish strong coupling between poly(A)-tail length and TE in HeLa cells. (**A**) Western blot showing siRNA-mediated depletion of PABPC1 and PABPC4 in HeLa cells. (**B**) The effect of depleting PABPC on coupling between tail length and TE in HeLa cells. Shown is the relationship between TE and median poly(A)-tail length after transfecting HeLa cells with the indicated siRNAs. A point representing the mRNA from one gene (*SPECC1*) fell outside the plot area in the siPABPC1&4 sample. Otherwise, this panel is as in Figure 2A.

**Figure supplement 5.**
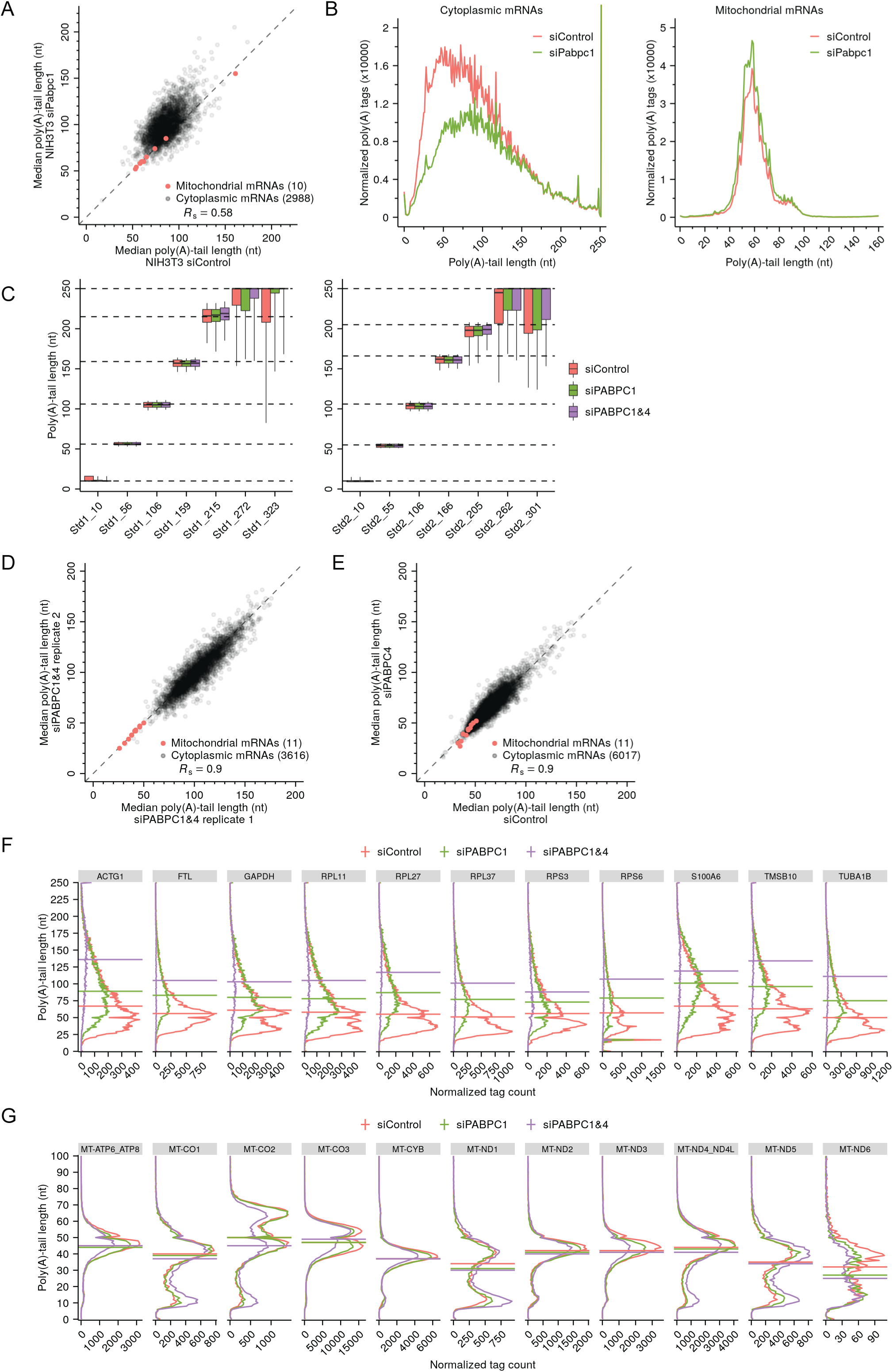
Support and extension of experiments showing that PABPC depletion causes premature decay of short-tailed mRNAs in HeLa cells. (**A**) The effect of PABPC knockdown on poly(A)-tail length in NIH3T3 cells. The plot compares median poly(A)-tail length in PABPC1-knockdown cells to that in control cells. Results are shown for mRNAs with ≥ 100 poly(A) tags (gray) and for mitochondrial mRNAs (red), merging data for *Mt-atp6* and *Mt-apt8*, and for *Mt-nd4* and *Mt-nd4l*, which are bicistronic mitochondrial mRNAs, and excluding data for *Mt-nd6*, which does not have a poly(A) tail. (**B**) The effect of PABPC knockdown on the abundance of mRNAs with different tail lengths in NIH3T3 cells. Shown are tail-length distributions of all cytoplasmic (left) and mitochondrial (right) mRNA poly(A) tags in control and PABPC1-knockdown cells. For each distribution, the abundance of tags was normalized to that of the spike-in tail-length standards. (**C**) The quality of the tail-length measurements, as determined using internal tail-length standards. Boxplots summarizing poly(A) tail-length distributions of two sets of tail-length standards spiked into total RNA samples obtained from HeLa cells transfected with the indicated siRNAs. Each box-whisker shows the 10^th^, 25^th^, 50^th^, 75^th^ and 90^th^ percentile of poly(A)-tail lengths. The dashed horizontal lines represent median tail lengths of individual standards, as determined by mobility in denaturing gels (Subtelny et al., 2014). (**D**) Reproducibility of tail-length measurements. The plot compares median poly(A)-tail lengths from two replicates of the double-knockdown HeLa cells, merging data for *MT-ATP6* and *MT-APT8*, and for *MT-ND4* and *MT-ND4L*, which are bicistronic mitochondrial mRNAs. (**E**) The effect of PABPC4 knockdown on poly(A)-tail length in HeLa cells. The plot compares median poly(A)-tail length in PABPC4-knockdown cells to that in control cells. Results are shown for mRNAs with ≥ 100 poly(A) tags (gray) and for mitochondrial mRNAs (red), merging data for *MT-ATP6* and *MT-APT8*, and for *MT-ND4* and *MT-ND4L*, which are bicistronic mitochondrial mRNAs. (**F**) The effect of PABPC knockdown on the tail-length distributions of top-expressed cytoplasmic mRNAs in HeLa cells. Plotted for mRNAs from each of the indicated genes are tail-length distributions in control, PABPC1- and double-knockdown cells, indicating median values with horizontal lines. For each distribution, the abundance of tags was normalized to that of the spike-in tail-length standards. (**G**) The effect of PABPC knockdown on the tail-length distributions of mitochondrial mRNAs in HeLa cells. Data were merged for *MT-ATP6* and *MT-APT8*, and for *MT-ND4* and *MT-ND4L*, which are bicistronic mitochondrial mRNAs. Otherwise, this panel is as in (**F**).

**Figure supplement 6.**
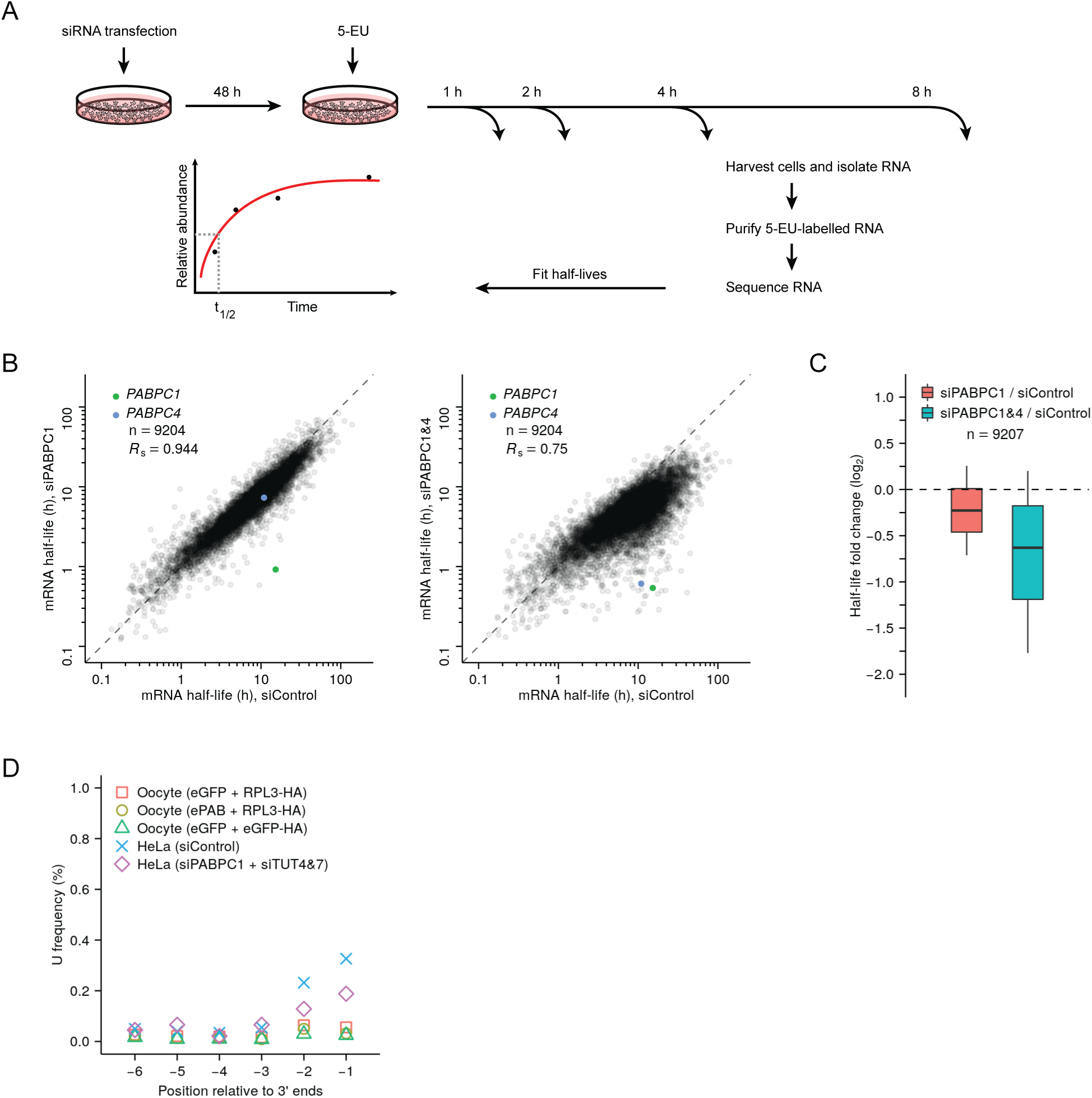
Measurements of mRNA half-lives in PABPC-depleted HeLa cells and terminal uridylation levels in frog oocytes. (**A**) Experimental scheme for measuring mRNA half-lives in HeLa cells. siRNA-transfected cells were incubated with 5-ethynyl uridine (5-EU), and cytoplasmic mRNA was harvested at the indicated time points. 5-EU-labeled RNA was biotinylated, isolated, and sequenced, using spike-in standards to normalize results from different time points. For labeled mRNA isolated from each gene, the approach to equilibrium was then fit to obtain its half-life. (**B**) The effect of PABPC knockdown on cytoplasmic mRNA half-lives in HeLa cells. For each gene, the mRNA half-life in PABPC1-knockdown cells (left) or in double-knockdown cells (right) is compared to that in control cells. Points representing mRNAs from three genes *RHOB*, *TSC22D3*, *PLEKHO2* fell outside the plot areas. (**C**) Distribution of the effects of PABPC knockdown on mRNA stability. Boxplots summarize the fold differences of mRNA half-lives observed in (**B**) when comparing PABPC1-knockdown or double-knockdown cells with control cells. Each box-whisker shows the 10^th^, 25^th^, 50^th^, 75^th^ and 90^th^ percentile. (**D**) Minimal terminal uridylation activity in frog oocytes. Plotted are the fractions of cytoplasmic mRNAs containing uridines near their termini, as detected in frog oocytes injected with mRNAs overexpressing the indicated proteins using tail-length sequencing. For comparison, results from HeLa cells transfected with the indicated siRNAs are replotted from Figure 5D.

**Figure supplement 7.**
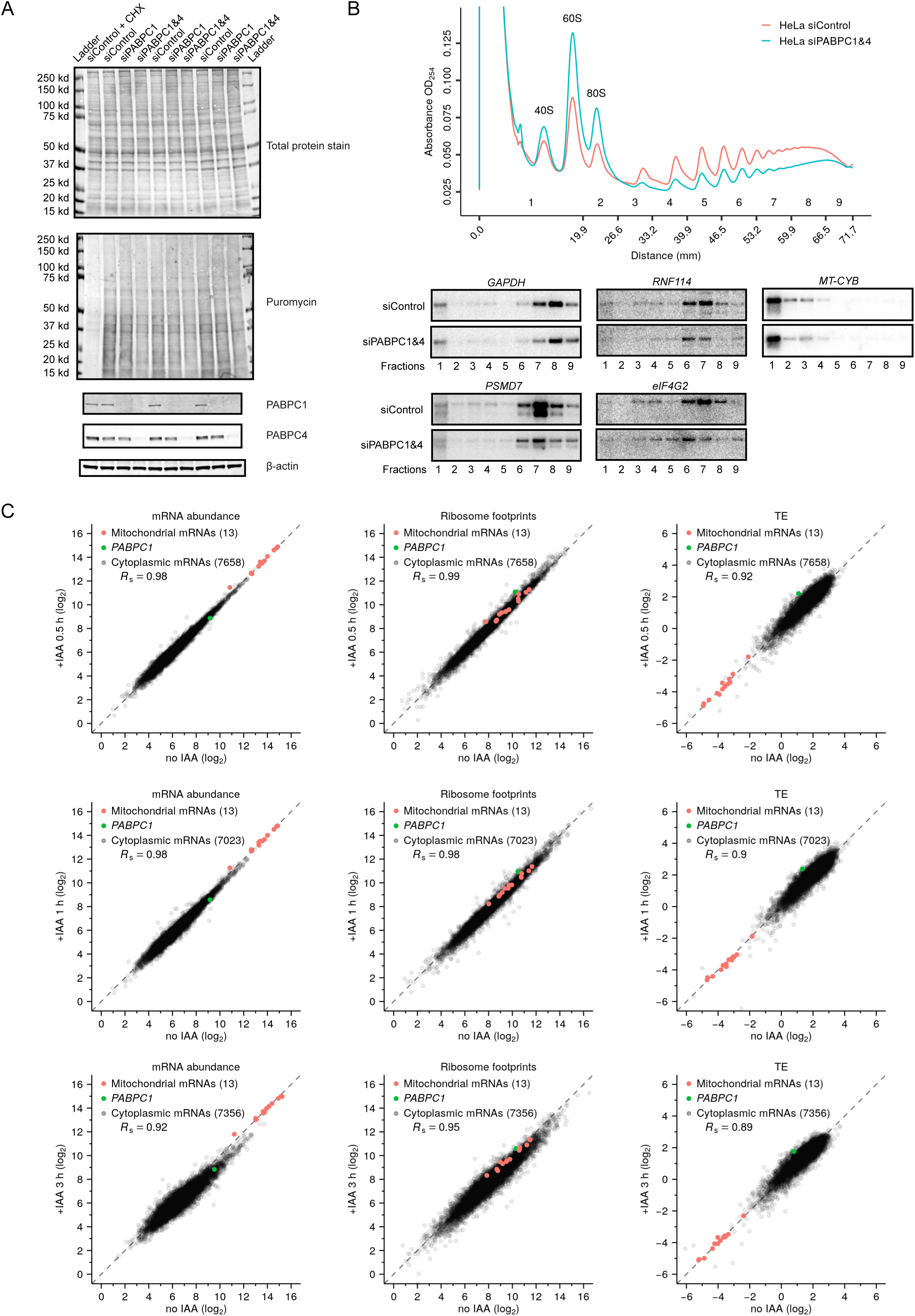
Depletion of PABPC in mammalian cell lines has minimal effect on TE. (**A**) The effect of PABPC knockdown on protein synthesis in HeLa cells. Cells transfected with the indicated siRNAs were cultured for 48 h, then treated with puromycin before harvesting for western-blot analysis. In a control sample, cycloheximide (CHX) was added prior to puromycin treatment. At the top is the membrane showing total protein levels. In the middle is the same membrane probed for puromycin to detect nascent protein synthesis. At the bottom are the results of the same membrane probed for the indicated proteins. (**B**) The effect of PABPC knockdown on polysomes in HeLa cells. At the top are polysome-gradient profiles from HeLa cells transfected with either control siRNAs or siRNAs targeting *PABPC1* and *PABPC4*. At the bottom are northern blots analyzing RNA collected from gradient fractions shown at the top, probing for four cytoplasmic mRNAs and one mitochondrial mRNA (*MT-CYB*). (**C**) The early effects of rapid PABPC1 degradation on mRNA abundance (left), ribosome-footprint abundance (middle) and TE (right) in HCT116 PABPC1-AID cells. At the top, values from cells treated with IAA for 0.5 h are compared to those from cells not treated with IAA. In the middle, values from cells treated with IAA for 1 h are compared to those from cells not treated with IAA. At the bottom, values from cells treated with IAA for 3 h are compared to those from cells not treated with IAA. For each gene, the number of mRNA and ribosome-footprint reads per kilobase are shown after normalizing to measurements for mitochondrial mRNAs. These results are summarized Figure 6F.

## Materials and methods

### Cloning and site-directed mutagenesis

All DNA plasmids were assembled by restriction-free cloning (Unger et al., 2010). Site-directed mutagenesis was also carried out with this method. For plasmids used for mammalian cell transfection, human PABPC1 (from HeLa cell cDNA) and *X. laevis* ePAB (from oocyte cDNA) coding sequences were cloned into pcDNA5/FRT/TO (Thermo Fisher, V652020). For siRNA-resistant human *PABPC1*, silent mutations were introduced at D107, K108, S109, I110, D111, N112, V131, C132, D249, E250, N252, and G253. Additional substitutions were made at I110L, D111E, D117E, A121G, G139A, Y140F, T147S, and R166K to disrupt the interaction between PABPC1 and eIF4G (Chorghade et al., 2017). For siRNA-resistant *X. laevis ePAB*, silent mutations were introduced at C128, K129, V130, V131, T249, E250, and N252. Sequences of oligos used for mutagenesis are provided in supplementary file 1. Plasmids and their sequence information will be available at Addgene.

### Templates for in vitro transcription

Plasmids for preparing DNA templates for in vitro transcription were assembled using the pGEM-11Zf(+) (Promega) backbone, inserting the appropriate sequence segments after the T7 promoter. HDV ribozyme sequence was obtained from the plasmid p2RZ (Avis et al., 2012). *X. laevis* PABPC1 (*pabpc1.S*), ePAB (*pabpc1l.L*) and RPL3 (*rpl3.L*) coding sequences were amplified from cDNA generated from *X. laevis* oocytes. *Renilla* (*Rluc*) and firefly luciferase (*Fluc*) coding sequences were obtained from pIS2 and pIS0, respectively (Farh et al., 2005). *NanoLuc* (*Nluc*) coding sequence was obtained from pNL1.1.TK (Promega). *X. laevis* b-globin 5′- and 3′-UTR sequences were obtained from pT7TS (Addgene #17091). Mouse *Malat1* 3′ sequence was obtained from the Comp.25 mutant plasmid (Wilusz et al., 2012). *Rluc* and *Fluc* reporters contained 5′- and 3′-UTR sequences inherited from the pGEM-11Zf(+) backbone, whereas *Nluc* reporters had the *X. laevis* b-globin 5′- and 3′-UTR sequences. Fragments containing variable poly(A) lengths were put in desired plasmids after all other DNA fragments were assembled, also using restriction-free cloning, except that C3040H competent cells (NEB) were used to amplify the assembled plasmids. Because long homopolymers tend to become shorter or get lost when plasmids are propagated in *E.coli*, individual clones were selected and checked by PCR and Sanger sequencing to confirm the desired length of each poly(A) region. These plasmid preparations were then used to generate templates for in vitro transcription without further propagation in *E.coli*.

For generating DNA templates for in vitro transcription, PCR reactions were carried out using KAPA HiFi HotStart PCR Kit, with a common 5′ primer upstream of the T7 promoter and different 3′ primers. For making RNAs ending in defined lengths of poly(A) sequence, a primer 300∼600 nt downstream of the HDV cleavage site was used to facilitate separating the 5′ cleavage product from the 3′ cleavage product and the uncleaved transcript. For making RNAs not ending in defined lengths of poly(A) sequence, a primer pairing to the end of the desired 3′ UTR sequence was used. All DNA templates for in vitro transcription were purified on agarose gels using the QIAquick Gel Extraction Kit (QIAGEN). Sequences of oligos used for generating DNA templates are provided in supplementary file 1.

### mRNAs for in vitro translation and oocyte injection

In vitro transcription was carried out with T7 RNA polymerase purified in house and used at 3.2 ng/µl final concentration in a standard 50 µl reaction containing 40 mM Tris pH 8.0, 21 mM MgCl_2_, 2 mM Spermidine (Sigma), 1mM dithiothreitol (DTT, GoldBio), 5mM NTP (buffered ATP, UTP, CTP, GTP mix, Thermo Fisher), 0.1 units yeast inorganic pyrophosphatase (NEB), 40 units SUPERase•In (Thermo Fisher), and 1µg DNA template. After incubation at 37°C for 2 h, 2 units of TURBO DNase (Thermo Fisher) were added, followed by another 20 min incubation at 37°C. For constructs with the HDV ribozyme, thermal cycling was performed to enhance HDV cleavage (65°C for 90 sec and 37°C for 5 min, 3 cycles, 50 µl of reaction per tube). Before gel loading, 2 µl 0.5M EDTA pH 8.0 and 50 µl 2x RNA Gel Loading Dye (Thermo Fisher, R0641) were added to all in vitro transcription reactions, regardless of whether the HDV cleavage step was performed or not. After incubation at 65°C for 5 min, RNAs were separated on 5% acrylamide denaturing gels (National Diagnostics, EC-829). Desired RNA bands were identified by UV-shadowing, excised, macerated and eluted in 300 mM NaCl at 23°C overnight on a rotator. The gels pieces were removed using Spin-X columns (Corning 8160), and RNAs were precipitated with isopropanol and resuspended in water for downstream reactions.

Capping of RNAs was carried out with the Vaccinia Capping System (NEB, M2080S) following the manufacturer’s protocol. RNAs were then purified by phenol/chloroform extraction and ethanol precipitation. Water-resuspended RNAs were applied to Micro Bio-Spin columns (Bio-Rad, 7326250) for desalting. All RNAs were checked for integrity by visualizing on formaldehyde-agarose denaturing gels before being stored at –80°C.

### *X. laevis* oocytes for in vitro translation and injection

*X. laevis* ovaries were obtained from Nasco (LM00935). Ovaries were broken down to 2–3 cm pieces with tweezers and incubated in calcium-omitted OR-2 buffer (82.5 mM NaCl, 2.5 mM KCl, 1 mM MgCl2, 1 mM Na2HPO4, 5 mM HEPES, pH 7.5) with 0.2% (w/v) collagenase A (Roche, 11088793001) on a rocker at 23°C for ∼3 h, at which point most oocytes were defolliculated. The oocytes were then washed extensively with calcium-omitted OR-2 buffer followed by complete OR-2 buffer (with 1 mM CaCl_2_). Stage V and VI oocytes were separated from the rest with a ∼0.8 mm diameter mesh sieve. For injection experiments, healthy stage VI oocytes were hand-picked and incubated in complete OR-2 buffer at 18°C overnight (> 16 h) for recovery before injection.

When preparing oocyte extracts, the bulk stage V and VI oocytes (500–1,000) were washed three times with ample Oocyte Extraction buffer (10 mM HEPES pH 7.5, 10 mM sodium acetate, 1 mM magnesium acetate and 2 mM DTT). After removing all excess buffer, oocytes were centrifuged at 20,000*g* for 20 min. The middle layer containing the extract was collected with a pipette and transferred to a new tube. The collected extract was centrifuged at 20,000*g* for 10 min, and the middle layer was again collected, avoiding the top lipid layer and the bottom insoluble layer as much as possible. The extract was passed through a 0.45 µm filter, then aliquoted and stored at –80°C. The concentration of the oocyte extract was measured using the Bradford assay (Thermo Fisher, 23236) and was usually at 35–45 mg/ml.

### In vitro translation

500 µl rabbit reticulocyte lysate obtained from Promega (L4151, untreated) was supplemented with 1 µl 12.5 mM Hemin (Sigma, 51280), 25 µl 1 M HEPES pH 7.5, and 2.5 µl 10 mg/ml yeast tRNA (Sigma, 10109495001). In vitro translation reactions were carried out with 5 ul of either supplemented rabbit reticulocyte lysate or oocyte extract in total volume of 20 µl containing 10 µM amino acid mix (Promega, L4461), 0.5 units SUPERase•In, 10 mM creatine phosphate (Roche 10621714001), 200 µg/ml creatine kinase (Roche, 10127566001), 20 mM HEPES pH 7.5, 75 mM potassium acetate, 1.5 mM magnesium acetate, and 1 fmol/µl *Rluc* or *Nluc* reporter RNA with indicated poly(A) tails. In addition, 2.5 fmol/µl *Fluc* RNA with a 120 nt poly(A) region followed by a mutant mouse *Malat1* 3′ end was included in each reaction for use as a normalization control. When examining the effect of adding purified proteins, reaction also included 2 µl either *X. laevis* PABPC1 (0.8 or 1.6 µg), *X. laevis* ePAB (0.8 or 1.6 µg), eGFP (1.6 µg) or buffer G (20 mM Tris pH 7.5, 150 mM NaCl, 5% glycerol, 5 mM DTT).

### Recombinant proteins

*X. laevis* PABPC1 (*pabpc1.S*) and ePAB (*pabpc1l.L*) coding sequences were amplified from cDNA generated from *X. laevis* oocytes. eGFP was obtained from the plasmid pCS2+-eGFP (Chen et al., 2017). All coding sequences were cloned into pET28a (Novagen) vector by restriction-free cloning. Each recombinant protein carried a hexahistidine tag at its C terminus. The plasmids were transformed into *E. coli* BL21(DE3) Star cells. After growing cells at 37°C to an optical density (OD_600_) of 0.6, expression of recombinant protein was induced with 0.5 mM isopropyl β-D-thiogalactoside (GoldBio). For purification of PABPC1 and ePAB, the cells continued to grow at 18°C for 16 h, after which they were collected by centrifugation, resuspended in 10 volumes of lysis buffer (20 mM Tris pH 7.5, 2 M NaCl, 5% (v/v) glycerol and 5 mM 2-mercaptoethanol) and lysed by sonication. Lysate was cleared by centrifugation at 25,000*g* for 30 min and incubated with Ni-NTA agarose (Qiagen, 30210) at 4°C for 1 h (0.5 ml resin per 50 ml supernatant). The resin was washed with 20 resin volumes of buffer H (20 mM Tris pH 7.5, 2 M NaCl, 5% glycerol, 5 mM 2-mercaptoethanol and 10 mM imidazole pH 7.5) and then with 20 resin volumes of buffer L (20 mM Tris pH 7.5, 150 mM NaCl, 5% glycerol, 5 mM 2-mercaptoethanol and 20 mM imidazole pH 7.5). Proteins were eluted with buffer E (20 mM Tris pH 7.5, 150 mM NaCl, 5% glycerol, 5 mM 2-mercaptoethanol and 200 mM imidazole pH 7.5). The eluate was diluted 1:9 with buffer CL (20 mM Tris pH 7.5, 25 mM NaCl, 5% glycerol, 5 mM DTT) and loaded onto a Mono S column (GE Healthcare). Bound proteins were eluted by linear NaCl gradient with buffer CL and buffer CH (20 mM Tris pH 7.5, 500 mM NaCl, 5% glycerol, 5 mM DTT). Fractions from the desired peak were pooled, concentrated with an Amicon filter unit (Millipore, UFC805024) and applied to a Superdex 200 column (GE Healthcare) in buffer G (20 mM Tris pH 7.5, 150 mM NaCl, 5% glycerol, 5 mM DTT). Fractions from the desired peak were pooled, concentrated with an Amicon filter unit, flash-frozen in liquid nitrogen and stored at −80°C. eGFP protein was purified similarly, except all buffers had 150 mM NaCl, and the cation-exchange step was omitted.

### Oocyte injections

Healthy *X. laevis* stage VI oocytes that had recovered from defolliculation were selected for injection. When examining the effects of expressing additional PABPC, either water or capped mRNA (2 pmol/µl) coding for either eGFP, PABPC1, or ePAB was injected into oocytes in a volume of 8 nl per oocyte (PLI-100 Plus Pico-Injector, Harvard Apparatus) at 23°C. Injected oocytes were incubated in complete OR-2 buffer at 23°C for 20 h, after which they were either lysed for making sequencing libraries or co-injected with a mixture of *Fluc* reporter mRNA (20 fmol/µl, with a 120 nt poly(A) region followed by a mutant mouse *Malat1* 3′ end) and either *Rluc* or *Nluc* reporter mRNA with indicated poly(A) tails (10 fmol/µl) in a total volume of 2 nl per oocyte, except in Figure 1F, where only 2 nl *Nluc* repoter mRNAs (10 fmol/µl) with indicated 3′ ends were injected. These oocytes were incubated in complete OR-2 buffer at 23°C for 6 h before being lysed for dual luciferase assays. When examining the effects of adding PABPC proteins, mixtures containing purified recombinant eGFP or ePAB protein (1.4 mg/ml), *Fluc* mRNA (20 fmol/ul), and short- or long-tailed *Rluc* reporter mRNA (10 fmol/ul) were injected into oocytes in a volume of 8 nl per oocyte. Dual luciferase assays were performed 6 hours after injections.

For PAL-TRAP, oocytes were injected with mRNA (1 pmol/ul) coding for either *X. laevis* RPL3-HA or eGFP-HA at a volume of 2 nl per oocyte. After incubation in complete OR-2 buffer at 23°C for one day, these oocytes were injected again with a mixture of mRNA coding for either eGFP or ePAB (2 pmol/ul) and a population of *Rluc* reporter mRNAs with different poly(A)-tail lengths (0, 30, 63, 98, and 120 nt, equimolar ratio; combined concentration, 166.5 fmol/ul) in a volume of 4 nl per oocyte. After incubation in complete OR-2 buffer at 23°C for another day, oocytes were collected and lysed for PAL-TRAP analysis.

### Luciferase assays

5–10 oocytes were lysed by vigorous shaking and pipetting in Passive Lysis Buffer (Promega, E1980), using 20 µl per oocyte. Lysates were cleared by centrifugation at 5,000*g* for 5 min at 4°C, after which 20 µl of each supernatant was transferred to a 96-well microplate. Luciferase assays were carried out with Dual-Luciferase Reporter Assay System (Promega, E1980) in a Veritas Microplate Luminometer according to the manufacturer’s protocol.

### Northern blots

Total RNA (1–10 µg) was separated on an agarose-formaldehyde gel (Mansour and Pestov, 2013) and transferred to a nylon membrane using the Whatman Nytran SuPerCharge (SPC) TurboBlotter system (Sigma, WHA10416300). After overnight transfer, RNA was crosslinked to the membrane using a UV Stratalinker 2400 at wavelength 254 nm for a total of 1200 µJ. For probing *Rluc* and 18S RNAs, membranes were pre-incubated with ULTRAhyb Ultrasensitive Hybridization Buffer (Thermo Fisher, AM8670) at 68°C under rotation for 1 h and then hybridized under the same conditions overnight with DNA probes, which were body-labeled with [⍺-^32^P]dCTP (PerkinElmer) using a Random Primer DNA Labeling Kit (TaKaRa, 6045) according to the manufacturer’s protocol. The templates for the labeling reactions were DNA fragments generated by PCR with gene-specific primers. After probe hybridization, membranes were washed two times (5 min each) with a low-stringency buffer (2X SSC and 0.1% SDS) at 68°C with rotation, and two times (15 min each) with high-stringency buffer (0.1X SSC and 0.1% SDS) at 68°C with rotation. For probing endogenous mRNAs, membranes were hybridized to radiolabeled gene-specific DNA oligonucleotide probes in ULTRAhyb-Oligo buffer (Thermo Fisher, AM8663) overnight at 42°C. The DNA probes were labeled with T4 PNK (NEB, M0201S) and [γ-^32^P]ATP (PerkinElmer). Following hybridization, membranes were washed three times (20 min each) with a wash buffer (2X SSC and 0.5% SDS) at 42°C with rotation. The blots were then analyzed using a Typhoon FLA 7000 phosphor-imager (GE Healthcare Life Sciences). Before probing for a second mRNA on the same blot, the blots were stripped three times (20 min each) in a boiling stripping buffer (0.04% SDS) with gentle shaking and then checked for any residual radioactivity by extended phosphorimaging. Sequences of oligos used for northern blots are provided in supplementary file 1.

### RNase H northern blots

Total RNA (1–10 µg) was mixed with a DNA oligo (100 pmol, supplementary file 1) complimentary to a segment of the 3′-UTR in water in a volume of 15 µl. In some experiments, two DNA oligos, each complementary to a different mRNA, were added together in one reaction. After incubating the RNA–DNA mix at 85°C for 5 min, the temperature was gradually lowered to 42°C (0.1°C per sec), and then 2 µl 10x Hybridase Buffer (500 mM HEPES pH 7.5, 1 M NaCl, 100 mM MgCl_2_), 0.5 µl Hybridase (Lucigen, H39500, 5 units/µl) and 2.5 µl water were added to the mix. After incubation at 42°C for 20 min, nucleic acids were extracted with phenol/chloroform, precipitated with ethanol, resuspended in 1x RNA Gel Loading Dye (Thermo Fisher, R0641), and resolved on an 8% acrylamide denaturing gel (National Diagnostics, EC-829). RNA was transferred onto Hybond-NX membranes (GE Healthcare, RPN303T) using a Trans-Blot SD Semi-Dry Transfer Cell (Bio-Rad). Membranes were incubated at 60°C for 1 h with EDC (1-ethyl-3-(3-dimethylaminopropyl)carbodiimide hydrochloride), Thermo Fisher, 22981) diluted in 0.17 M 1-methylimidazole pH 8 to chemically crosslink 5′ phosphates to the membrane. Blots were hybridized to radiolabeled DNA oligonucleotide probes in ULTRAhyb-Oligo buffer (Thermo Fisher, AM8663) overnight at 42°C. The DNA probes were complimentary to the 3′-UTR regions immediately adjacent to the poly(A) tails and were labeled with T4 PNK (NEB, M0201S) and [γ-^32^P]ATP (PerkinElmer). Following hybridization, membranes were washed three times (20 min each) with a wash buffer (2X SSC and 0.5% SDS) at 42°C with rotation. The blots were then analyzed using a Typhoon FLA 7000 phosphor-imager. Before probing for a second RNA on the same blot, the blots were stripped as described for conventional northern blots. Sequences of oligos used for RNase H northern blots are provided in supplementary file 1.

### Western blots

Samples for western blots were boiled in NuPAGE LDS Sample Buffer (Thermo Fisher, NP0007), resolved on NuPAGE 4-12% Bis-Tris protein gels (Thermo Fisher, NP0323BOX), and transferred to 0.2 µm PVDF membranes (Thermo Fisher, LC2002) with a Mini Gel Tank (Thermo Fisher, A25977), according to the manufacturer’s protocol. Membranes were then blocked with 5% (w/v) non-fat dry milk in TBS buffer (20 mM Tris pH 7.5, 150 mM NaCl) for 30 min. Primary antibodies were diluted at 1:1,000 in TBST buffer (20 mM Tris pH 7.5, 150 mM NaCl, 0.1% (v/v) Tween 20) and incubated with the blot at 4°C overnight. Secondary antibodies were diluted at 1:10,000 in TBST buffer and incubated with the blot at 23°C for 1 h. Blots were analyzed using an Odyssey Clx machine (LI-COR) and the Image Studio software (LI-COR). For total protein detection, Revert 700 Total Protein Stain Kits (LI-COR, 926-11010) were used before incubation with primary antibodies, according to the manufacturer’s protocol. Total protein levels were quantified with the ImageQuant TL software (GE Healthcare Life Sciences).

### ^35^S Labeling in oocytes

Groups of five oocytes were incubated with 0.5 mCi/ml EasyTag EXPRESS ^35^S protein labeling mix (PerkinElmer, NEG772007MC) in 100 µl complete OR-2 buffer at 23°C for 1 hour. For a control, a group of oocytes was pretreated with 100 µg/ml cycloheximide in 100 µl complete OR-2 buffer for 5 min before adding the labeling mix. After labeling, buffers were removed and oocytes were washed with 1 ml complete OR-2 buffer. After removing all residual wash buffer, 100 µl ice cold lysis buffer (20 mM HEPES pH 7.5, 100 mM KCl, 5 mM MgCl_2_, 1% (v/v) Triton X-100, 1x Halt proteinase inhibitor cocktail (Thermo Fisher, 78429), 2 mM DTT) was added to each group, and oocytes were lysed by vigorous shaking and pipetting. Lysates were cleared by centrifuging at 5,000*g* at 4°C for 5 min, and supernatants were boiled in NuPAGE LDS Sample Buffer and resolved by SDS-PAGE. Gels were dried and analyzed using the Typhoon FLA 7000 phosphor-imager and the ImageQuant TL software (GE Healthcare Life Sciences).

### PAL-TRAP

Injected oocytes (∼100) were washed once with complete OR-2 buffer and three times with 1 ml ice-cold buffer RL (20 mM HEPES pH 7.5, 100 mM KCl, 5 mM MgCl_2_, 1% (v/v) Triton X-100, 100 µg/ml cycloheximide, 20 units/ml SUPERase•In, cOmplete protease inhibitor cocktail (Sigma, 11836170001, 1 tablet per 10 ml buffer)). After removing all wash buffer, oocytes were lysed in buffer RL (10 µl/oocyte) by vigorous shaking and pipetting. Lysates were cleared by centrifugation at 5,000*g* at 4°C for 10 min. A portion of each supernatant (5%) was saved as the input for western analysis and RNA isolation. The remaining supernatant was incubated with anti-HA magnetic beads (Thermo Fisher, 88837) using 5 µl slurry per 100 µl supernatant. A population of *Nluc* RNAs with varied poly(A)-tail lengths (29, 63, and 139 nt, equimolar ratio) were spiked in at 5 ng per 100 µl supernatant, and the bead mixture was incubated at 4°C for 1 h with end-to-end rotation. Beads were then immobilized using a magnetic stand, and the supernatant was removed, with a portion (5%) saved as the flowthrough for western blot and RNA isolation. Beads were washed four times with 1 ml buffer RL and resuspended in 110 µl buffer RL, 10 µl of which was taken for western blot and the remainder was used for RNA isolation. RNA was isolated with 10 volumes of Tri Reagent (Thermo Fisher, AM9738) according to the manufacturer’s protocol. 1µg purified RNA was incubated at 37°C for 30 min with 10 units of T4 PNK (NEB, M0201S) in PNK buffer containing 20 units SUPERase•In in a 25 µl reaction to remove 2′-3′-cyclic phosphates of reporter RNAs. Two PAL-seq v3 libraries were made from each input and eluate sample, and after monitoring reproducibility, data from each pair of replicates were merged during analyses, whereas only one library was made from the each flowthrough sample.

### Polysome gradients

Lysate preparation and centrifugation were performed the same as for ribosome profiling, except that no nuclease was added prior to centrifugation. RNA was purified using 10 volumes of Trizol LS (Thermo Fisher, 10296010) according to manufacturer’s protocol and proteins were extracted by chloroform-methanol precipitation.

### Cell culture

All mammalian cells were cultured at 37°C with 5% CO_2_. HeLa cells were cultured in DMEM (VWR, 45000-304) with 10% FBS (TaKaRa, 631106). HCT116 OsTIR1 cells (kindly provided by Masato Kanemaki) and their derivatives were cultured in McCoy’s 5A media (Thermo Fisher, 16600082) supplemented with 10% FBS, 2mM L-glutamine (Thermo Fisher, 25030081). NIH3T3 cells were cultured in DMEM with 10% BCS (Sigma, 12133C). HCT116 PABPC1-AID cells were cultured as HCT116 OsTIR1 cells but supplemented with 600 µg/ml G418 (Thermo Fisher, 10131027). ZF4 cells were obtained from ATCC and cultured in DMEM F-12 media (ATCC, 30-2066) and 10% FBS, at 28°C with 5% CO_2_.

### Transfection

For siRNA transfection, cells were plated at 2 x 10^6^ cells per 10 cm plate, cultured for 12 h, and transfected with 15 pmol total siRNAs and 45 µl Lipofectamine RNAiMAX (Thermo Fisher, 13778500) in 1 ml Opti-MEM media (Thermo Fisher, 31985062) according to manufacturer’s protocol. Cells were split 1:4 onto new plates 24 h after transfection and then collected for analysis 48 h (Figure 5–6, Figure 5—figure supplement 6B–C and Figure 6—figure supplement 7) or 72 h (Figure 4, Figure 4—figure supplement 4–5 and Figure 5—figure supplement 6D) after transfection. For rescue of PABPC-knockdown, cells were plated at 0.6 x 10^6^ cells per well in 6-well plates, and transfected with 2.5 pmol total siRNAs and 7.5 µl Lipofectamine RNAiMAX in 250 µl Opti-MEM media. After 24 h, cells were transfected with 1 µg DNA and 3 µl FuGENE HD (Promega, E2311) in 150 µl Opti-MEM media according to manufacturer’s protocol, and collected for analysis 40 h after DNA transfection. For other DNA plasmid transfections, cells were plated at 0.6 x 10^6^ cells per well in 6-well plates, cultured for 24 h, transfected with 1 µg DNA and 3 µl FuGENE HD (Promega, E2311) in 150 µl Opti-MEM media according to manufacturer’s protocol. siRNAs were all purchased from Dharmacon, including siControl (D-001810-10-05), human siPABPC1 (L-019598-00-0005), human siPABPC4 (L-011528-01-0005) and mouse siPabpc1 (L-060385-00-0005). Plasmids and their sequence information will be available at Addgene.

### Puromycin-based translation assay

Assays were performed as described (Schmidt et al., 2009). Puromycin (Thermo Fisher, A1113802) was added to cell culture media at final concentration 1 µg/ml. Cells were incubated at 37°C for 10 min before harvest. For a control, cycloheximide was added to the media at final concentration 100 µg/ml 5 min before the addition of puromycin.

### IAA-induced PABPC1 degradation

AID was introduced at the C terminus of PABPC1 using Cas9-mediated genome engineering in HCT116 OsTIR cells. The 5′ and 3′ homology arms (∼500 nt) flanking the stop codon of PABPC1 were amplified from genomic DNA of HCT116 OsTIR1 cells and cloned into a donor plasmid (kindly provided by Iain Cheeseman). The AID coding sequence followed by a the T2A peptide and mCherry, whose sequences were obtained from pMK1221 (Addgene, #84220), was inserted immediately before the PABPC1 stop codon. The PAM region targeted by Cas9 on the donor plasmid was mutated. The Cas9 guide RNA was cloned into pX330-BFP, as described (McKinley and Cheeseman, 2014). Both the donor and the Cas9 plasmids were co-transfected into HCT116 OsTIR cells at 0.5 µg each in a 6-well format. After 72 h, single cells with strong mCherry signal were sorted with flow cytometry into 96-well plates. Clones were expanded and genotyped by PCR and western blot. Only clones that were homozygous for AID integration were retained. One clonal line (sC152-C16) was used for results shown in Figure 5C. 6 h after siRNA transfection, 1 µg/ml doxycycline was added to induce *OsTIR1*. After another 18 h, indole-3-acetic acid (IAA, GoldBio) was dissolved in ethanol and added to cells at a concentration of 0.5 mM. 24 h after IAA addition, cells were harvested for analysis. Sequences of oligos used for making the donor and the Cas9 plasmids are provided in supplementary file 1.

IAA-induced depletion of PABPC1 in the PABPC1-AID cell line (sC152-C16) was relatively slow and generally required more than 12 h for 90% depletion. To achieve faster depletion dynamics, more copies of *OsTIR1* were introduced. *OsTIR1* coding sequences were obtained from pBabe Puro osTIR1-9Myc (Addgene #80074) and subcloned into PiggyBac-dCas9-Tet1 (Liu et al., 2016) (kindly provided by Shawn Liu), replacing the dCas9 coding sequence. This plasmid was co-transfected with piggyBac Transposase expression vector (System Biosciences, PB210PA-1) into the PABPC1-AID cell line (sC152-C16). 48 h after transfection, cells were selected in 600 µg/ml G418 for 10 d. Single clones were expanded and checked for *OsTIR1* expression. The clone with the highest *OsTIR1* expression (sC278-C2) was picked for IAA-induced PABPC1-AID degradation. To minimize background degradation, cells were incubated with 1 µg/ml doxycycline and 0.2 mM auxinole (Aobious, AOB8812), an inhibitor of OsTIR1 (Yesbolatova et al., 2019) for 6 h, after which fresh media with 1 µg/ml doxycycline and 0.5 mM IAA were added. Cells were collected after 0.5, 1, and 3 h of induction. For the non-induced condition, fresh media with 1 µg/ml doxycycline but no IAA were added, and cells were collected after 1 h. Plasmids used for making the PABPC1-AID cell lines and their sequence information will be available at Addgene.

### Ribosome profiling and matched RNA-seq

To prepare lysate from mammalian cells, cycloheximide (CHX) was added to each plate at final concentration 100 µg/ml. Plates were immediately moved to a cold room and culture media were removed. Cells were washed twice with ice-cold PBS supplemented with 100 µg/ml CHX. After the last wash, 1 ml buffer RLL (20 mM HEPES pH 7.5, 100 mM KCl, 5 mM MgCl_2_, 1% (v/v) Triton X-100, 100 µg/ml cycloheximide, 500 unit/ml RNasin Plus (Promega, N2615), 2 mM DTT, cOmplete protease inhibitor cocktail (Sigma, 11836170001, 1 tablet per 10ml buffer)) was added to each 15 cm plate, and cells were scraped off and incubated on ice for 10 min. The resulting lysates were passed through a 26-gauge needle six times and cleared by centrifugation at 1300*g* at 4°C for 10 min. To prepare lysate from frog oocytes, injected oocytes were washed once with complete OR-2 buffer and three times with buffer RLL. After removing all wash buffer, oocytes were lysed in buffer RLL (10 µl/oocyte) by vigorous shaking and pipetting. Lysates were cleared by centrifugation at 5,000*g* at 4°C for 10 min. One tenth of each cleared lysate was added to 10 volumes of Tri Reagent for total-RNA preparation, following the manufacturer’s protocol. Another small portion (∼10 µl) was taken for western analysis. The remainder was aliquoted, flash frozen in liquid nitrogen, and stored at –80°C.

For ribosome profiling, RNase I (Thermo Fisher, AM2294) was added to lysate, using 30 units per OD_260_ unit for lysate from mammalian cells and 10 units per OD_260_ unit for lysate from frog oocytes. After incubation at 23°C for 30 min, lysates were loaded onto a 10–50% sucrose gradient (20 mM HEPES pH 7.5, 100 mM KCl, 5 mM MgCl_2_, 10–50% (w/v) sucrose, 100 µg/ml cycloheximide, 20 units/ml SUPERase•In, 2 mM DTT), centrifuged in a Beckman ultracentrifuge with SW41Ti rotor at 36,000 rpm for 2 h, and then fractioned on a BioComp gradient fractionator. Fractions corresponding to 80S ribosomes were collected, except in experiments that generated data for Figure 6A and B, in which fractions corresponding to 40S, 60S, and 80S ribosomes were collected so as to avoid loss of the smaller-sized mitochondrial ribosomes (Rooijers et al., 2013). Collected fractions were concentrated with an Amicon filter unit (Millipore, UFC810024), and ribosome-protected RNA fragments were released in a buffer containing 20 mM HEPES pH 7.5, 100 mM KCl, 5 mM MgCl_2_, 2 mM EDTA, 20 units/ml SUPERase•In, 2 mM DTT. Released RNA fragments were incubated with 0.2 mg/ml Proteinase K (Thermo Fisher, AM2548) in the presence of 1% SDS at 42°C for 20 min, and then purified by phenol/chloroform extraction and isopropanol precipitation.

Sequencing libraries were prepared as described previously (Subtelny et al., 2014), except in experiments that generated data for Figure 6A and B, in which different size markers were used to isolate RNA fragments from a larger size range (20–40 nt in the first gel) so as to avoid loss of mitochondrial ribosome-protected fragments (Rooijers et al., 2013).

For matched RNA-seq, rRNAs were either depleted with the RiboZero Gold Kit (Illumina, Figures 2A, Figure 2—figure supplement 2A–B, and Figure 4—figure supplement 4B), depleted with the NEBNext rRNA Depletion Kit (Human/Mouse/Rat, NEB, Figures 6 and S6), or not depleted (Figures 2B, 2E, 2F, and Figure 2—figure supplement 2C). Because the RiboZero Gold Kit depletes mitochondrial mRNAs (Figure 2—figure supplement 2C), mitochondrial mRNAs were not used for normalization of data generated with this kit. Total RNA (with rRNA depleted or not depleted) was fragmented by incubating at 95°C for 20 min in RNA Fragmentation buffer (2 mM EDTA, 12 mM Na_2_CO_3_, 88 mM NaHCO_3_) and then ethanol precipitated. RNA fragments were size-selected and sequenced in parallel with ribosome-protected fragments. A detailed protocol for ribosome profiling is available at http://bartellab.wi.mit.edu/protocols.html.

Sequencing was performed on an Illumina HiSeq 2500, with standard runs of either 40 or 50 cycles. Reads were trimmed at both ends to removed adapter sequences using Cutadapt (v1.18) (Martin, 2011) and then mapped to their respective genome reference using STAR (v2.4.2a) with the parameters “--runMode alignReads --outFilterMultimapNmax 1 -- outReadsUnmapped Fastx --outFilterType BySJout --outSAMattributes All --outSAMtype BAM Unsorted SortedByCoordinate”. Mapped reads were counted for each gene with htseq-count (0.11.0). For both ribosome profiling and RNA-seq, only reads that uniquely mapped to coding regions of annotated genes (excluding the first 15 codons and the last 5 codons) were included in downstream analyses. For TE analyses, an expression cutoff of 30 RNA-seq reads was applied for each gene, with no cutoff for ribosome-footprint reads. Normalizations of RNA-seq and ribosome-footprint reads were performed with DESeq2 (Love et al., 2014), considering reads for all genes passing the cutoff, except when absolute TE comparisons were made between two samples—in which case, RNA-seq and ribosome-footprints data were normalized by only considering reads from mitochondrial genes using software DESeq2 (Love et al., 2014). When only relative TEs were considered, TEs were manually centered at 0.

### TAIL-seq

Our implementation of TAIL-seq (Eisen et al., 2020) resembled that of mTAIL-seq (Lim et al., 2016) in that it used splint ligation to append the 3′ adapter, as in PAL-seq v1 (Subtelny et al., 2014). Total RNA (1–30 µg) was mixed with two sets of tail-length standards (0.1 ng per 1 µg total RNA for each set) (Subtelny et al., 2014), and trace 5′-radiolabeled marker RNAs (Eisen et al., 2020), which were used to evaluate tail-length measurements and 3′ ligation efficiency, respectively. These RNAs were ligated to a 3′ adapter in a 64 µl reaction containing 1.5 µM 3′ adapter, 1.25 µM 3′ splint oligo, 0.2 mM ATP, 1 unit/µl RNasin Plus (Promega, N2615), 10 mM MgCl_2_, 1x T4 RNA Ligase 2 reaction buffer (NEB, M0239S), and 0.5 units/µl T4 RNA Ligase 2 (NEB, M0239S), with the enzyme added after the other components had been mixed and incubated at 23°C for 5 min. The ligation reaction was incubated at 18°C for 18 h. Small portions (2 µl) were removed at the start and end of the reaction for examining the ligation efficiency. After ligation, RNA was extracted with phenol/chloroform, precipitated with ethanol, resuspended in 10 µl water, and mixed with 90 µl 1x RNA Sequencing Buffer from the RNase T1 kit (Thermo Fisher, AM2283). After incubation at 50°C for 5 min and on ice for 5 min, 1 µl RNase T1 (1 unit/µl, Thermo Fisher, AM2283) was added, and the reaction was incubated at 23°C for 15 min, followed by phenol/chloroform extraction and precipitation with the Precipitation/Inactivation Buffer in the RNase T1 kit. RNA was resuspended in 12 µl RNA Gel Loading Dye and RNA fragments ranging between 100 and 760 nt were purified on an 8% acrylamide denaturing gel, captured on streptavidin beads, 5′ phosphorylated, and ligated to an equal mix of four phased 5′ adapters as described (Eisen et al., 2020). cDNA was generated using SuperScript III (Thermo Fisher, 18080044) with DNA primers containing barcodes used for multiplexing, eluted from beads by base hydrolysis of RNA and resolved on 6% urea-acrylamide denaturing gels as described (Subtelny et al., 2014), except that cDNA fragments with sizes between 150 nt and 760 nt were selected. cDNAs were sequenced directly using a HiSeq 2500 machine in a paired-end 50-by-250 run in normal mode with a v3 kit as described (Eisen et al., 2020). Poly(A)-tail lengths shown in Figures 2, Figure 2—figure supplement 2, Figure 4, Figure 4—figure supplement 4 and Figure 5—figure supplement 5C–G were measured with this method. Sequences of adapters and other oligos used for TAIL-seq are provided in supplementary file 1.

### PAL-seq v3

PAL-seq v3 differs from PAL-seq v2 (Eisen et al., 2020) in two major ways: 1) Barcodes were embedded in the 3′ adapters so that different samples could be pooled for downstream stages of library construction. 2) Sequencing was performed with cDNA rather than PCR-amplifed cDNA. Each reaction was set up in a total volume of 20 µl by mixing total RNA (0.3–3 µg) with two sets of tail-length standards (0.1 ng of each set per 1 µg total RNA) (Subtelny et al., 2014), two splint DNA oligos (55 pmol A-splint and 2.75 pmol U-splint, which allowed polyadenylated RNAs with a terminal uridine to be more efficiently captured) and one barcoded 3′ adapter (50 pmol). Poly(A)-selected mRNA from zebrafish ZF4 cell line (0.1 ng per ug total RNA) was also added to mammalian RNA samples to enable additional assessment of tail-length measurement reproducibility. The RNA–DNA mix was incubated at 65°C for 5 min, and the temperature was gradually lowered to 16°C (0.1°C per sec) before other component were added to a volume of 30 µl containing 0.2 mM ATP, 1 unit/µl RNasin Plus, 10 mM MgCl_2_, 1x T4 RNA Ligase 2 reaction buffer and 0.5 units/µl T4 RNA Ligase 2. After incubation at 16°C for 16 h, EDTA (1 µl, 0.5 M, pH 8.0) was added to stop ligation and samples to be sequenced together (with different barcodes) were combined, phenol/chloroform extracted, and precipitated with ethanol. Ligated RNA was resuspended in 20 µl water, mixed with 180 µl RNase T1 kit buffer (20 mM sodium citrate pH 5.0, 1 mM EDTA, 7 M urea), heated to 50°C for 5 min, and chilled on ice for 5 min, before adding 1.6 µl RNase T1 (1 unit/µl, Thermo Fisher, AM2283). After incubating the reaction at 23°C for 30 min, RNA was extracted with phenol/chloroform and precipitated using the Precipitation/Inactivation Buffer in the RNase T1 kit. Fragments between 150 and 760 nt were gel-purified, selected on beads, 5′-end phosphorylated, ligated to a 5′ adapter, and used for cDNA synthesis in the same way as in our implementation of TAIL-seq, except the 5′ adapter lacked the barcodes used for multiplex sequencing and cDNAs with lengths between 200 and 760 nt were selected. Sequencing was performed similarly as in PAL-seq v2 (Eisen et al., 2020) but with some differences: 1) 1 fmol cDNA rather than PCR-amplified cDNA was used per lane. 2) A single primer was used to obtain two reads. 3) After cluster generation and sequencing-primer hybridization, and before extension of the primer through the poly(A)-tail region using the Klenow fragment, 16 dark cycles were performed in order to extend the sequencing primer past the barcode and constant regions of the 3′ adapters as well as past two nucleotides corresponding to the RNA 3′ termini. 4) 40 cycles of standard sequencing-by-synthesis were performed to yield the first sequencing read (read 1). 5) After obtaining read 1, the flow cell was stripped, the same sequencing primer was annealed, and 260 cycles of standard sequencing-by-synthesis were performed to read the barcode (6 nt, which required 7 cycles), a constant segment of the 3′ adapter (8 nt, which required only 7 cycles because of an extra cycle in the barcode region), and the sequence of the RNA, beginning at its 3′ terminus, which revealed whether the RNA had a terminal uridine and provided information used to measure the length of the poly(A) tail. Poly(A)-tail lengths shown in Figure 3, Figure 3—figure supplement 3, Figure 4—figure supplement 5A–B and Figure 5 were measured with this method. Sequences of adapters and other oligos used for PAL-Seq v3 are provided in supplementary file 1.

### PAL-seq v4

For PAL-seq v4, several changes were made from PAL-seq v3. The 3′-ligation reaction was essentially the same, except that one 3′ adapter was used for all samples. After the ligation, each sample was processed separately rather than mixed. Ligated RNA was resuspended in 10 µl water and fragmented in a 100 µl rather than 200 µl volume, and fragments with sizes between 130 and 760 nt were then gel purified. For reverse transcription, different primers containing the multiplexing barcode sequences as well as a region with 5 random nucleotides was used. After cDNA elution from the beads, half of each sample was saved at –20°C for future use. cDNA samples to be sequenced on the same lane in a flow cell were combined, ethanol-precipitated and gel-purified as in PAL-seq v3, except cDNA fragments with sizes between 190 nt and 800 nt were selected. Sequencing cluster generation was performed similarly to PAL-seq v3, using 0.3 fmol cDNA mixture for each lane. Read 1 started with 12 cycles of standard sequencing-by-synthesis that first sequenced the 5-nt random region (used to call clusters) and then sequenced the 6-nt barcode region (which required 7 cycles) with a custom primer. The flow cell was stripped, a second sequencing primer annealed, and 2 dark cycles were performed in order to extend this primer past the two nucleotides corresponding to the RNA 3′ termini. The custom extension of the primer through the poly(A) tail region with Klenow was then performed, as in PAL-seq v2 (Eisen et al., 2020) and v3, followed by 40 cycles of standard sequencing-by-synthesis to complete the read 1, which was generated using two sequencing primers and had a total length of 52 nt (12 cycles before and 40 cycles after the Klenow reaction). The flow cell was then stripped, and the second sequencing primer was used for 255 cycles of standard sequencing-by-synthesis to generate read 2. Poly(A) tail lengths shown in Figure 6E were measured with this method. Sequences of adapters and other oligos used for PAL-Seq v4 are provided in supplementary file 1.

### Genome references and gene annotations

Both human (release 25, GRCh38.p7, primary assembly) and mouse (release 10, GRCh38.p4, primary assembly) genomic sequences were downloaded from the GENCODE website. Sequences of mitochondrial pseudo genes (supplementary file 2) in human and mouse genomes were masked to avoid losing mitochondrial mRNA reads due to multi-mapping. *X. laevis* genomic sequences were downloaded from the Xenbase website (v9.1 assembly, repeat masked), and scaffolds were removed so that only chromosomal sequences would be considered. The *X. laevis* mitochondrial genomic sequence was obtained from NCBI website (NC_001573.1) and appended to the *X. laevis* genome.

Both human (release 25, GRCh38.p7, main annotation) and mouse (release 10, GRCh38.p4, primary assembly) gene annotations were downloaded from the GENCODE website. *X. laevis* gene annotations (v9.1 assembly, v1.8.3.2 primary transcript) were downloaded from the Xenbase website, and only chromosomal annotations were used. *X. laevis* mitochondrial gene annotations were curated based on the information obtained from the NCBI website (NC_001573.1) and appended. For all species, annotations for protein-coding genes were extracted, and for each gene, the isoform with the longest open reading frame (ORF) was selected to represent that gene. In cases in which multiple isoforms had ORFs of the same length, the isoform with the longest transcript was selected as the reference annotation. For TAIL-seq and PAL-seq analyses, the databases were supplemented with sequences and annotations of tail-length standards, *eGFP*, *Rluc*, *Fluc*, and *Nluc*. For ribosome profiling and matched RNA-seq analyses, the databases were supplemented with coding sequences and annotations of *eGFP*, *Rluc*, *Fluc*, and *Nluc*. For RNA-seq analyses used to determine half-lives, the databases were supplemented with coding sequences and annotations of *AcGFP* (Eisen et al., 2020), *Rluc*, and *Fluc*. All supplemented sequences are provided in supplementary file 2.

### Annotation of mRNA 3′-end isoforms

Each PAL-seq tag corresponding to an mRNA provides the site of cleavage and polyadenylation, which enabled annotation of mRNA 3′-end isoforms. Uniquely mapped tags from all PAL-seq datasets in the same cell type (either HeLa or frog oocyte) were merged. RNA-seq reads generated for comparison to ribosome profiling were similarly merged. Software HOMER (Heinz et al., 2010) was used to call peaks by using the merged RNA-seq data as the background and the merged PAL-seq data as the signal, with the following parameters “-style factor -o auto - strand separate -fdr 0.001 -ntagThreshold 50 -fragLength 40 -size 40 -inputFragLength 30 - center”. The peaks that intersected with annotated protein exons were retained as unique mRNA 3′ ends.

### TAIL-seq data analysis

Phased constant sequences at the 5′ end and poly(A) sequences at the 3′ end of read 1 were removed with a custom script. The trimmed sequences were mapped to reference genomes with STAR (v2.4.2a) with the parameters “--runMode alignReads --outFilterMultimapNmax 1 -- outReadsUnmapped Fastx --outFilterType BySJout --outSAMattributes All --outSAMtype BAM Unsorted SortedByCoordinate”. Mapped sequences were intersected with sequences of protein-coding genes by bedtools (v2.26.0) with the parameters “intersect -wa -wb -bed -s”, retaining only those sequences that were assigned to a single gene. Remaining clusters were then filtered, requiring at least 5 combined N and T bases in the first 6 nucleotides of read 2. For each library, 1% of the filtered read clusters (but no more than 50,000 and no less than 5,000) were randomly picked as the training set for determining the Hidden Markov Model.

For each Illumina sequencing cluster, average intensities of each channel from position 15–50 of read 1 were used to normalize intensities of each channel in read 2. Then a T-signal value for each cycle of read 2 in each cluster was calculated by dividing normalized intensity from T channel by the sum of normalized intensities from the other three channels. If the T-signal was 0 for a specific cycle, the T-signal values from neighboring cycles (up to 10, minimum 5) were averaged to infer the value for that cycle. If a cluster had more than 5 cycles with a read 2 T-signal value of 0, the cluster was discarded. A five-state mixed Gaussian Hidden Markov Model (from python ghmm package) was then used to decode the sequence of states that occurred in read 2. It consisted of an initiation state (state 0), a strong poly(A)-tail state (state 1), a weak poly(A)-tail state (state 2), a weak non-poly(A)-tail state (state 3) and a strong non-poly(A)-tail state (state 4). All reads started in state 0, and all states were only allowed to go forward (from 0 to 4). The model was initialized with the following transition probability matrix (from state in row to state in column):

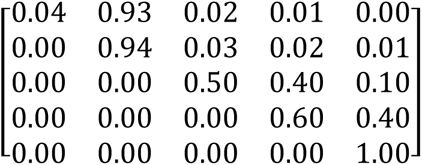

The emission matrix for the mixed population 1 was initialized with (states in row, mean, variance and population fraction in column):

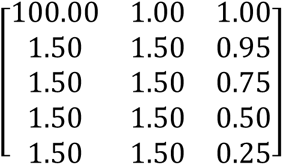

The emission matrix for the mixed population 2 was initialized with (states in row, mean, variance and population fraction in column):

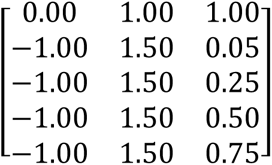

After model initialization, all clusters from the training set were used to perform unsupervised training, and then the trained model was used to decode the sequence of states for all retained clusters. For each cluster, the poly(A) tail length was determined by summing the number of states in state 1 and 2. Each cluster assigned to a specific gene by read 1 was considered as a poly(A) tag. When evaluating tail-length distributions of mRNAs from individual genes, only results from genes with at least 100 tags were considered. Note that the HiSeq 2500 machine in high-throughput mode raises the laser intensities after 50 cycles in read 2, causing irregular T-signal at this position and lower-than-expected state transition predicted by the Hidden Markov Model. This led to a mild depletion of poly(A) tags with tail lengths called at 50 nt, but it did not affect results and conclusions made from overall tail-length distributions.

For normalization of poly(A) tags among samples, DESeq2 (Love et al., 2014) was used with all spike-in tail-length standards to obtain the scaling factor for each dataset. When analyzing median tail lengths, only genes with poly(A) tag counts exceeding an indicated cutoff were included in the analyses.

### PAL-seq data analysis

For PAL-seq v3, read 1 sequences were mapped to reference genomes with STAR (v2.4.2a) with the parameters “--outFilterMultimapNmax 1 --outFilterType BySJout --outSAMattributes All -- outSAMtype BAM SortedByCoordinate”. Mapped sequences were intersected with annotations for protein-coding genes by bedtools (v2.26.0) with the parameters “intersect -wa -wb -bed -S”, retaining only those tags for which the read 1 sequence was assigned to a single gene or to an mRNA isoform, when mRNA 3′-end annotations were used. Remaining clusters were then filtered by first removing the first 14 bases of read 2 (which corresponded to a constant region and the barcode region) and then requiring at least 5 combined N and T bases within the next 6 nucleotides of read 2. (This filtering step was skipped for analyses of terminal uridylation in Figure 5). For each library, 1% of the filtered read clusters (but no more than 50,000 and no less than 5,000) were randomly picked as the training set for determining the Hidden Markov Model. The sequence of poly(A) states was determined similarly as for TAIL-seq, except a single Gaussian Hidden Markov Model was used, and the emission matrix was initialized with (states in row, mean and variance in column):

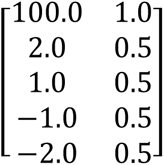

For normalization of poly(A) tags among samples, DESeq2 (Love et al., 2014) was used with all spike-in tail-length standards to obtain the scaling factor for each dataset. When analyzing median tail lengths, only mRNA isoforms with poly(A) tag counts exceeding an indicated cutoff were included in the analyses. Data analyses for PAL-seq v4 were the same as for PAL-seq v3, except before mapping, the first 12 nt of read 1 were trimmed to remove the random region and the barcode region, and no nucleotides of read 2 were trimmed. Note that the HiSeq 2500 machine in rapid-run mode raises the laser intensities after 101 cycles in read 2 of a PAL-seq v4 run, causing irregular T-signal at this position and lower-than-expected state transition predicted by the Hidden Markov Model. This led to a mild depletion of poly(A) tags with tail lengths called at 101 nt, but it did not affect results and conclusions made from overall tail-length distributions.

### PAL-seq analysis of terminal uridylation

A variant HeLa cell line was used for data shown in Figures 5D–F and Figure 5—figure supplement 6D. This Flp-In T-Rex HeLa cell line (Thermo Fisher, discontinued) had an *RPL3-HA* gene that had been inserted using the Flp-In (Thermo Fisher) system. We have no reason to suspect that the unique features of this line affected any results shown. Data were processed as in PAL-seq v3, except that mapped reads were intersected with HeLa mRNA 3′-end annotations (see Annotation of mRNA 3′-end isoforms). All poly(A) tags with poly(A)-tail lengths ≥ 2 nt were used when examining the presence of U nucleotides content near the ends of poly(A) tails.

### Half-life measurements

48 h after siRNA transfection, HeLa cells were incubated with pre-warmed fresh media with 400 µM 5-ethynyl uridine (5EU, Jena Biosciences) for 1, 2, 4, and 8 h. Cells were removed from plates by treatment with trypsin, washed once with ice cold PBS and lysed with 200 µl ice cold buffer RL. Lysates were cleared by centrifuging at 1300*g* for 5 min. Supernatants were transferred to 2 ml Tri Reagent, and total RNA was prepared according to the manufacturer’s protocol.

Biotinylation of 5EU-labeled RNA and purification of biotinylated RNA were performed at described (Eisen et al., 2020). RNA-seq libraries were prepared from purified 5EU-labeled RNA and from total RNA with the NEXTflex Rapid Directional mRNA-seq Kit (Bioo Scientific, 5138-10). Sequencing was performed on an Illumina HiSeq 2500 with a standard 40-cycle run. Sequencing reads were mapped with STAR (v2.4.2a) with parameters “--runMode alignReads -- outFilterMultimapNmax 1 --outReadsUnmapped Fastx --outFilterType BySJout -- outSAMattributes All --outSAMtype BAM Unsorted SortedByCoordinate”. Exon-mapped reads were counted for each gene with htseq-count (0.11.0).

Relative 5EU-labeled mRNA levels for each gene at each time point were obtained by normalizing read counts based on the counts for a 5EU-containing *GFP* RNA standard that had been spiked into each sample prior to 5EU biotinylation (Eisen et al., 2020). The steady-state mRNA levels of each gene were measured as the average of normalized read counts obtained from sequencing the input RNA for the 1 and 8 h time points. 987 genes with values that differed by > 2-fold at these two time points were excluded from analysis. The nls package in R was used to fit the following equation for each gene:

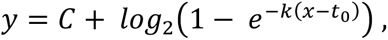

where *x* is the labelling time in hours (using 9999 h as the labeling time for the steady-state data point), *y* is log_2_ value of the normalized read count (level of labeled RNA), and *t*_0_ is the time offset. *k* is the decay constant, which was used to determine the half-life *t*_1/2_ using

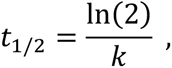

and *C* is a coefficient determined by both *k* and the synthesis rate *S*, such that

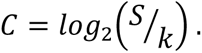

To fit *t*_0_ for each condition, the value of *t*_0_ was varied from 0.05 to 0.7, with an interval of 0.05, and the value that gave the smallest mean square-loss of *y* when fitting to data from all genes was used. When fitting to results for each gene, *C* was bound by (0, Inf), and *k* was bound by (0, Inf). *C* was initialized with *max*(*y*), and *k* was initialized with 0.23. Genes without a converged fit (27 of 9644, 17 of 9646, and 12 of 9728 in analysis of the siControl, siPABPC1, and siPABPC1&4 samples, respectively) were omitted from down-stream analyses.

### Statistical analysis

Graphs were generated and statistical analyses were performed using R (R Core Team, 2017). Statistical parameters including the value of n, statistical test, and statistical significance (*P* value) are reported in the figures or their legends. No statistical methods were used to predetermine sample size. Statistical tests for correlations (between nonoverlapping dependent groups) were performed based on a published method (Silver et al., 2004) using R package cocor (Diedenhofen and Musch, 2015). For luciferase assays of injected oocytes, each replicate refers to a group of 7–10 oocytes. For ^35^S labeling of injected oocytes, each replicate refers to a group of 5∼8 oocytes. For the puromycin-based translation assay, each replicate refers to a separate transfection experiment.

